# Dormancy, not apoptosis, restricts hematopoietic stem cell mutagenesis during aging

**DOI:** 10.64898/2026.05.09.724021

**Authors:** Foteini Fotopoulou, Megan Druce, Ruzhica Bogeska, Charles Imbusch, Viktoria Flore, Melanie Ball, Julia Knoch, Susanne Lux, Esther Rodríguez-Correa, Kleo Aurich, Ian Ghezzi, Marleen Büchler-Schäff, Ana-Matea Mikecin, Jeyan Jayarajan, Daniel B. Lipka, Adrian Baez-Ortega, Inigo Martincorena, Benedikt Brors, Alex Cagan, Michael D. Milsom

## Abstract

Genome instability and mutagenesis are hallmarks of aging, acting as drivers of some age-associated pathologies, including cancer^1–3^. Somatic cells engage multiple layers of protection against mutagenesis, including detoxification of genotoxic metabolites; repair of DNA damage; and elimination of cells which suffer excessive damage^4–7^. In this context, the intrinsic apoptotic pathway is engaged in response to activation of the DNA damage response (DDR) and is thought to play a major role in limiting accumulation of mutations, particularly in cells that act as an origin for cancer, such as somatic stem cells^8,9^. However, the dissection of the relative contribution of different protective mechanisms that restrict mutagenesis in such cells is confounded by the long time frame of experiments; relatively low mutation burden in non-malignant cells; and high variance across individuals due to differences in germ line and environment. Here we employ deep whole-genome sequencing (WGS) combined with extended time-course sampling from a range of experimental mouse models to study mutation acquisition in hematopoietic stem cells (HSCs) during aging. Having validated that murine HSCs recapitulate mutation acquisition patterns observed in aged human HSCs, we made the surprising discovery that apoptosis has a negligible role in restricting mutagenesis. Instead, we found that HSC dormancy inhibits mutagenesis during normal aging, with dormant HSCs from old mice demonstrating a mutation burden akin to their young counterparts. Importantly, breaking HSC dormancy via induction of sterile inflammation led to a dramatic acceleration in mutation rate, demonstrating that non-genotoxic environmental stimuli can modulate genome stability. These findings provide new insights into the correlation between inflammation and both aging and carcinogenesis.

## Introduction

At a fundamental level, the generation of DNA mutations requires a chemical alteration in the molecular structure of DNA, typically as a result of damage, followed by the conversion into a new stable structure, often as a consequence of errors in DNA repair. The cell employs a multi-layered array of mechanisms in order to maintain genome stability, such as engaging metabolic pathways which reduce the production of endogenous genotoxic compounds, or which detoxify both endogenous and exogenous DNA-damaging agents^10^. Downstream of genotoxic damage, the DNA damage response (DDR) coordinates a spectrum of partially redundant DNA repair pathways in order to try and resolve the many differing forms of DNA damage that a cell can encounter^11^. However, the high levels of damage that a cell is exposed to on a daily basis^3^ ultimately results in these protective mechanisms reducing mutation acquisition rates, as opposed to an outright prevention of mutagenesis. In this context, the cell can also restrict the propagation of mutated DNA by initiating programmed cell death via the process of cell intrinsic apoptosis, also known as the mitochondrial pathway. The DDR can directly trigger the intrinsic apoptotic pathway once a certain threshold of signalling has been achieved, which acts to maintain the balance between elimination of cells with high levels of DNA damage versus the excessive destruction of cells with only minor damage, which would likely compromise tissue function^12^. Thus, the intrinsic apoptotic pathway is thought to act as a critical gatekeeper which restricts the rate of mutation acquisition with the passage of time.

DNA damage and the resulting genomic instability that this provokes are thought to play a central role in promoting in the ageing process, contributing to age associated tissue degeneration and dysfunction, as well as the evolution of age associated diseases such as cancer^1–3^. Within regenerative tissues such as the hematopoietic system, it is of particular interest to understand the processes that regulate mutagenesis within the stem cell compartment during aging. This is because these cells: (i) are very long-lived and can therefore accumulate many mutations during aging; (ii) continually propagate these mutations to all their differentiated progeny, potentially compromising tissue function; and (iii) act as a likely cell of origin for cancer, which is ultimately driven by the accumulation of cooperating mutations within a single clone^9^. While retrospective analysis of mutational patterns can provide some indication of the processes which promote DNA mutation in stem cells during normal aging, cause-and-effect relationships between prospective *in vivo* modulators of genome stability and their mutational consequences can only be firmly established using an interventional approach using experimental model systems. In this study, we employ the murine hematopoietic system as a model tissue to study the mechanisms that govern the quantitative and qualitative acquisition of mutations within the stem cell compartment during physiologic aging, in the absence of exposure to frank DNA damaging agents such as chemotherapy or ionizing irradiation.

### Accurate and sensitive quantification of clonal mutation burden in HSCs

To accurately quantify the genome-wide mutation burden within individual long-term HSCs (LT-HSCs) from inbred C57BL/6J mice, we adopted the previously documented approach of expanding single purified LT-HSCs *in vitro*, followed by whole genome sequencing (WGS) of DNA isolated from the resulting clonal progeny^13^ (**Fig. S1**, **Fig S2a**). Germline variants were determined by whole genome sequencing of paired DNA from tails and were then subtracted from mutations called in the LT-HSC-derived colony. To remove mutations acquired during the *in vitro* expansion process, variants with an allele frequency (VAF) lower than 0.3 were excluded. In contrast to studies in which somatic point mutations have been identified and subsequently used as spontaneously occurring barcodes to deduce the ancestorial relationships between different HSC clones, our goal was to generate an accurate and sensitive quantitative and qualitative assessment of the genome-wide mutation spectrum of each HSC clone. Since the per genome estimate of mutation burden can be affected by insufficient sequencing depth^14^, we initially performed a benchmarking exercise to establish the necessary coverage required to accurately call single nucleotide variants (SNVs) and insertion-deletions (indels). To these ends, an individual LT-HSC-derived colony and its corresponding germline were sequenced to a coverage of 89X and 97X, respectively. Alignments of both sets of WGS data were then randomly downsampled in triplicate, in decremental steps of 10X coverage. SNVs and indels were separately called for each downsampled dataset using either CaVEMan^15^ or Pindel^16^ and the SNVs/indels which co-occurred in both the 89X sample and each downsampled data set were annotated as “confident calls” (**Fig S2b**). We identified that for both SNVs and indels, the absolute number of variants called and the number of confident calls began to plateau at around 30X coverage, capturing approximately 87% of confident SNVs and 72% of confident indels (**Fig. S2c-j**). We therefore set 30X as our minimum threshold for coverage and normalized per-genome mutation estimates based on the proportion of the mappable genome sequenced in each sample, which is a higher coverage than that normally used to identify SNVs for the purpose of clonal barcoding and hierarchical clustering.

### Mutational patterns acquired in murine LT-HSCs during aging are analogous to those observed in HSCs from elderly humans

We next applied this approach to characterize the process of mutation acquisition during normal aging. Individual LT-HSCs were isolated from young adult (8 months old) and healthy old mice (24 months old), and the mutation burden was determined following *in vitro* clonal expansion (**Fig. 1a**). The frequency of LT-HSCs which could form colonies was equivalent between young and old donors, suggesting no evidence for sample bias during *in vitro* expansion due to phenomena such as differing levels of senescence in the old and young HSC clones (**Fig 1b**). In line with previous studies^17^, old LT-HSCs demonstrated an approximate two-fold higher level of SNVs compared to young HSCs, with an average acquisition rate of 44 mutations per genome per year during normal aging (**Fig 1c-d**). In contrast, the frequency of indels and structural variants (SVs) per LT-HSC genome was much lower and only showed a non-significant increase with age (**Fig 1e-g**, **Fig. S3a-b**). As expected, given the largely neutral accumulation of somatic mutations, the vast majority of detected variants occurred at non-coding regions of the genome, with no age-dependent shift in this distribution (**Fig. S3c**). A more detailed analysis of mutation patterns can be achieved by the analysis of the sequence context surrounding the site of a given point mutation, using SNV trinucleotide spectra^18^. This analysis revealed that previously defined mutational patterns identified in aged human HSCs were also prevalent in aged murine LT-HSCs, including single base substitution (SBS) signatures “HSPC”, “SBS1”, “SBS5” and “SBS18”^17,19–21^ (**Fig. 1h**, **S4a-c**). We additionally estimated the number of SNVs in each clone that belonged to each of these mutational signatures, demonstrating a significant age-associated increase in both HSPC- and SBS5-attributed mutations in LT-HSCs (**Fig. 1i-n**). Taken together, these data suggest that somatic mutagenesis in murine HSCs is dominated by the same mutational processes as those acting in human HSCs during normal aging.

**Figure 1.**
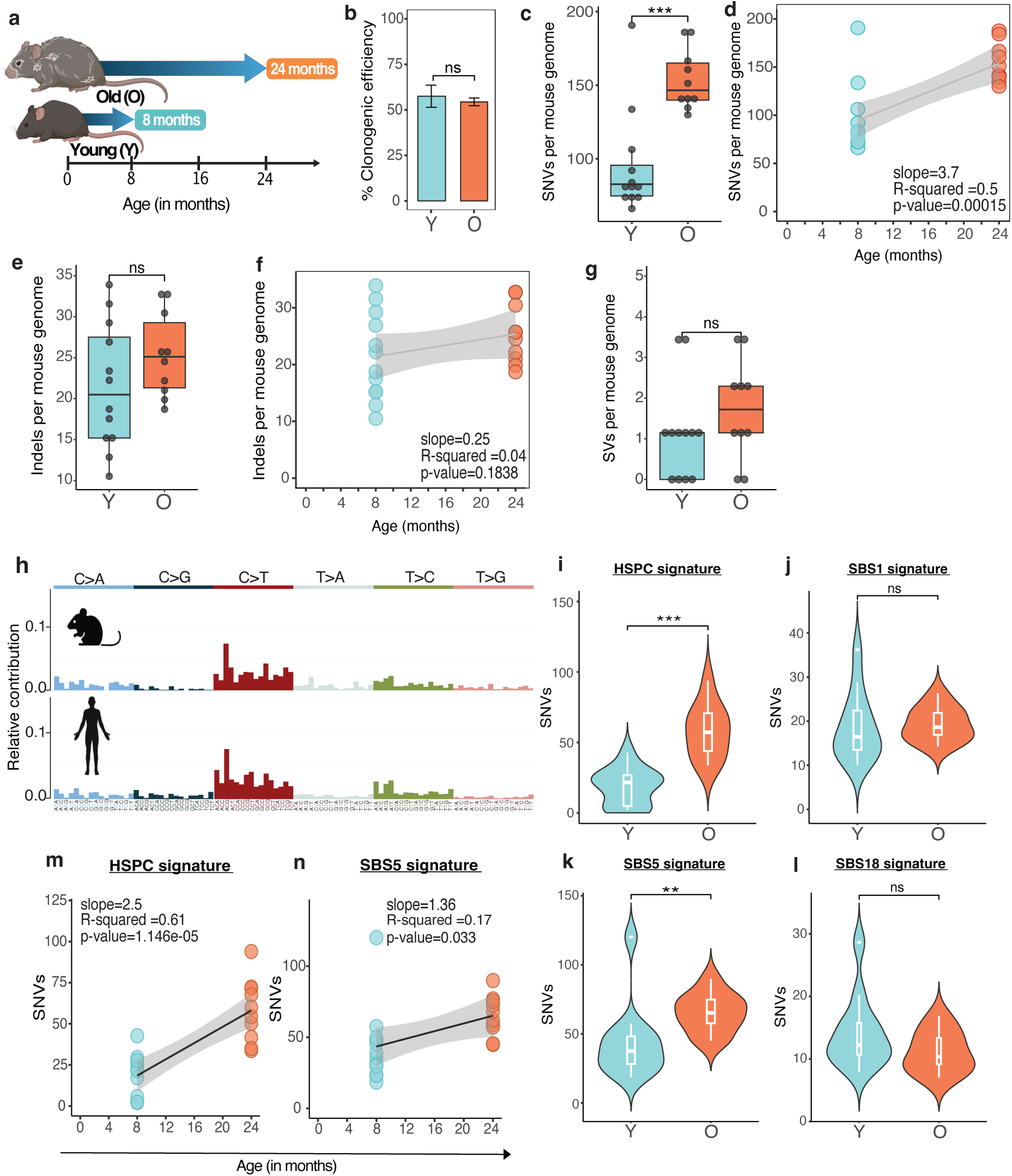
Endogenous age-associated somatic mutagenesis in murine HSCs. **a)** Schematic overview of experimental design. LT-HSCs were isolated from young (8 months) and old (24 months with no obvious pathology) C57BL/6J mice and expanded in culture, prior to WGS. **b)** Clonogenic efficiency per plate, defined as the percentage of single-sorted LT-HSCs that divided at least once (>=2 cells/colony). 192 colonies were seeded (2 plates) per mouse, for n=3 mice per group. The bars indicate the mean +/- SEM. **c)** Absolute number of SNVs per LT-HSC, normalized according to the proportion of the callable genome for each sample. **d)** Linear regression of SNV burden as a function of age, with 95% confidence intervals (CI) indicated by shading. **e)** Absolute number of indels per LT-HSC, normalized as in **c**. **f**) Linear regression model estimating the rate of indel accumulation per month. **g)** Absolute number of SVs per LT-HSC, normalized as in **c**. **h)** Trinucleotide SNV spectra of murine and human LT-HSCs. The murine spectrum was generated using all SNVs from old LT-HSC clones. The human spectrum is based on substitution data from a healthy donor (HD1957) reported by Abascal *et al.*^21^ **i-l)** Absolute number of SNVs attributed to the indicated fitted signature after bootstrapping. **m-n)** Linear models imposed on the signature-specific burdens from **i** or **k**, as indicated, estimating the rate of signature-specific mutation accumulation over time. 95% CI is indicated. ns: p>0.05, **: p<0.01, ***: p<0.001, using the Mann-Whitney U test. LT-HSC were isolated from 3 different mice for each group, with each dot representing a single LT-HSC. Y: young (n=12), O: old (n=10). Box plots indicate median, inter-quartile range (IQR) with whiskers extending to 1.5x IQR.

### Cell intrinsic apoptosis does not restrict mutagenesis in the HSC pool during normal aging

The induction of programmed cell death via the cell intrinsic apoptosis pathway is a well-characterized consequence of both DNA damage and defects in downstream DNA repair^11,22,23^. Cell intrinsic apoptosis has long been regarded as a key mechanism which ensures genomic stability of HSCs, with some suggestions that HSCs have a lower threshold for inducing apoptosis in order to protect against mutagenesis in a potential cancer cell of origin^24–28^. In order to formally interrogate the role of apoptosis in restricting HSC mutagenesis *in vivo* during normal physiologic aging, we made use of an established genetic mouse model which disables the intrinsic apoptotic pathway by preventing the key signalling step of release of cytochrome c into the cytoplasm via conditional genetic ablation of the Bak Bax pore in the mitochondrial membrane^29–31^. Thus, the *Bak*^-/-^ *Bax*^fl/fl^ alleles were combined with the *Scl-Cre-ERT* transgene^32^ and the resulting mice were subject to tamoxifen treatment at 8-weeks of age in order to delete *Bax* in LT-HSCs that already harboured a homozygous deletion of *Bak* (**Fig. 2a**). LT-HSCs were subsequently purified from these mice at 8-months of age, and their clonal mutation burden was determined following *in vitro* expansion, as described above. 87% of expanded LT-HSC clones demonstrated homozygous deletion of *Bax* (*Bax*^Δ/Δ^) and were selected for further analysis (**Fig. 2b**). Surprisingly, this analysis demonstrated that there was no quantitative difference in the per genome levels of either SNVs, indels nor SVs within *Bak*^-/-^ *Bax*^Δ/Δ^ LT-HSCs compared to LT-HSCs isolated from age-matched wild-type (WT) mice (**Fig. 2c-e**), suggesting that the loss of apoptotic function did not alter the mutation acquisition rate under normal physiologic conditions, across a window of time where WT LT-HSCs would be predicted to gain more than 20 SNVs (**Fig. 1c**). In line with this observation, a qualitative analysis of SNV mutation patterns revealed that *Bak*^-/-^ *Bax*^Δ/Δ^ LT-HSCs demonstrated analogous SBS signatures to those observed in WT LT-HSCs, dominated by the typical C to T transitions that occur during normal aging (**Fig. 2f**). Probabilistic attribution of per cell levels of SBS-associated mutations demonstrated no significant differences in levels of HSPC, SBS5 and SBS18-derived mutations compared to WT LT-HSCs (**Fig. 2g-j**). Thus, the overall mutational profiles of WT and *Bak*^-/-^ *Bax*^Δ/Δ^ HSCs remained highly similar, although we did detect a modest decrease in absolute levels of SBS1-derived mutations in *Bak*^-/-^ *Bax*^Δ/Δ^ LT-HSCs (**Fig. 2g-k**, **Fig. S4d**). Taken together, these data indicate that cell intrinsic apoptosis plays no major role in restricting mutation acquisition in LT-HSCs *in vivo* under physiologic conditions and suggest that the processes underlying the progressive acquisition of mutations during normal aging are not normally sufficient to trigger programmed cell death.

**Figure 2.**
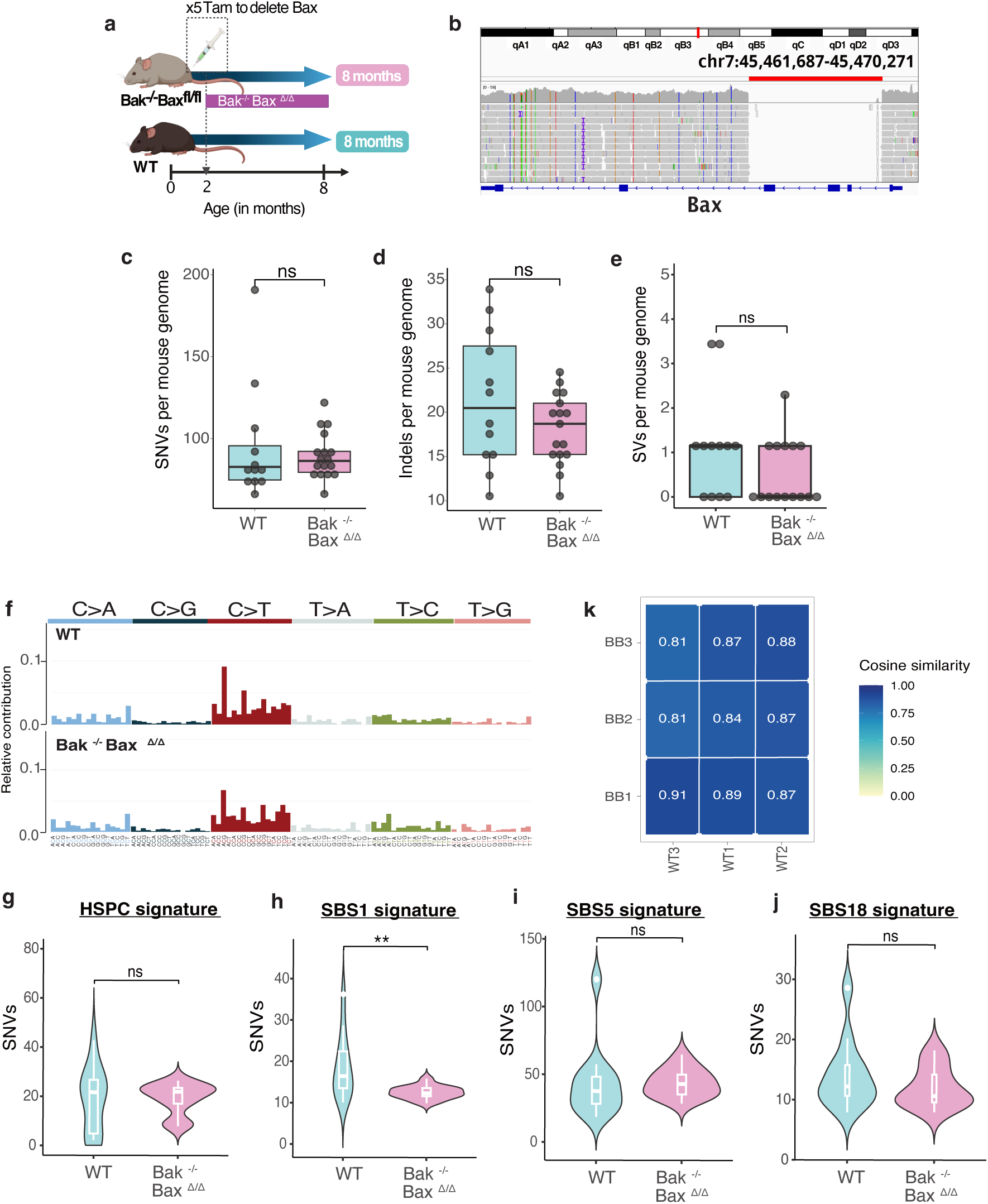
Genetic abrogation of cell intrinsic apoptosis does not impact on mutagenesis in LT-HSCs under physiologic conditions *in vivo*. **a)** Schematic overview of the experimental groups. *Scl-Cre-ERT2 Bak*^-/-^ *Bax*^fl/fl^ mice were treated with tamoxifen at 2 months of age to induce the *Bax* deletion in LT-HSCs. Post-recombination, the mice are referred to as *Bak*^-/-^ *Bax*^Δ/Δ^. LT-HSC were harvested from wild type (WT) and Bak^-/-^ Bax^Δ/Δ^ mice at 8 months of age, expanded clonally *in vitro,* then subject to WGS as in Fig. 1. **b)** Integrative Genome Viewer (IGV) visualization of successful Cre-induced homozygous recombination at the *Bax* locus (chr7:45,461,687-45,470,271) in a representative LT-HSC-derived colony. The red bar indicates the deleted region spanning exons 2-4. The coverage drop in the highlighted region reflects loss of reads mapping to the excised sequence. A representative subset of the total sequenced reads for this region is displayed. **c-e)** Absolute number of **c)** SNVs; **d)** indels; and **e)** SVs, normalized according to percentage coverage of the mouse genome. Each dot represents a single LT-HSC. **f)** Trinucleotide SNV spectra of WT HSCs (top) compared to *Bak*^-/-^ *Bax*^Δ/Δ^ HSCs (bottom). SNVs have been merged across all LT-HSC colonies by group. **g-j)** Absolute number of SNVs attributed to the indicated fitted signature. **k)** Heatmap showing the cosine similarity of SNV trinucleotide spectra between WT and *Bak*^-/-^ *Bax*^Δ/Δ^ (BB) HSCs grouped per mouse (1-3). All pairwise comparisons were evaluated using the Mann-Whitney U test. ns: p>0.05, **: p<0.01. LT-HSCs were isolated from 3-5 different mice, with n=12 (WT) and n=17 (Bak^-/-^ Bax^Δ/Δ^) clones analysed per group. The mouse IDs BB3 corresponds to three different, but co-housed, mice that were merged and collectively analysed as a single mouse for more robust signature analysis. Each dot represents a single LT-HSC. Box plots indicate median, inter-quartile range (IQR) with whiskers extending to 1.5x IQR.

### Maintenance of a long-term dormant state restricts mutagenesis in LT-HSCs during aging

We and others have characterized that a subset of LT-HSCs possess a low proliferation history throughout the majority of a mouse’s life span, by sustaining cell cycle quiescence for extended periods of time^33,34^. This state of so-called dormancy appears to protect HSCs from acquiring features of aging, and correlates with very low levels of DNA damage^33–35^. One might therefore speculate that dormancy might protect LT-HSCs from mutation acquisition during aging. However, it has also been proposed that HSCs are particularly vulnerable to the accumulation of DNA damage during quiescence^36,37^, while the observation in humans of near-identical mutation burdens in HSCs and peripheral granulocytes from age-matched donors, despite their different proliferative histories, suggests that HSCs may acquire mutations through processes that are independent of cell division^21^. In order to directly interrogate the mutational burden of dormant LT-HSCs versus their more actively cycling counterparts, we made use of a second genetic mouse model which allows sub-segregation of LT-HSCs based on their divisional history *in vivo*. Thus, the LT-HSCs within the *Scl-tTA H2B-GFP* mouse model accumulate the H2B-GFP fusion protein within chromatin, until the expression of the label is switched off by supplying doxycycline in the drinking water. The label is then subsequently diluted as the LT-HSCs divide, allowing the prospective identification and purification of dormant label retaining cells using flow cytometry^38^. We treated a cohort of *Scl-tTA H2B-GFP* mice with doxycycline for a duration of 18 months, starting at 4 months of age (**Fig. 3a**). As previously documented^34^, it was possible to discern LT-HSCs in 22-month-old mice which had retained label above background levels throughout the lifetime of the mouse, allowing us to interrogate the mutation burden of these dormant cells relative to more actively cycling LT-HSCs from the same animals (**Fig. 3b**). Despite being the same chronologic age, dormant LT-HSCs demonstrated a significantly lower level of SNVs per genome than their non-label retaining counterparts and showed a trend towards lower levels of indels (**Fig 3c-d**). Linear regression analysis indicated that dormant LT-HSCs accumulated SNVs at a rate that was approximately three-fold slower than active HSCs during physiologic aging (**Fig. 3e**). A more detailed analysis of mutation patterns between dormant and active LT-HSCs from the same aged mice revealed that there was no difference in the genomic regions in which mutations occurred, nor in the type of indels that we observed (**Fig. S5a-c**). A comparison of SNV mutation signatures demonstrated a broadly similar mutation profile between dormant and active LT-HSCs (**Fig. 3f-j**, **Fig.S5d**). However, the absolute levels of mutations corresponding to the SBS1 “clock-like” signature, which has been associated with cell division^39^, were significantly enriched in active LT-HSCs, which cross-validates the model’s capacity to prospectively isolate LT-HSCs with a lower proliferation history (**Fig. 3h**). Interestingly, dormant LT-HSCs also demonstrated significantly lower levels of the SBS18 signature which is attributed to DNA damage from reactive oxygen species (ROS)^40^, correlating with our previous work which demonstrated that the dormant state protected LT-HSCs from genotoxic damage from metabolic ROS *in vivo*^35^ (**Fig. j**). Overall, the data from this model indicates that LT-HSCs accumulate mutations in a heterogeneous fashion during normal aging, with dormant LT-HSCs being more resistant to mutation acquisition.

**Figure 3.**
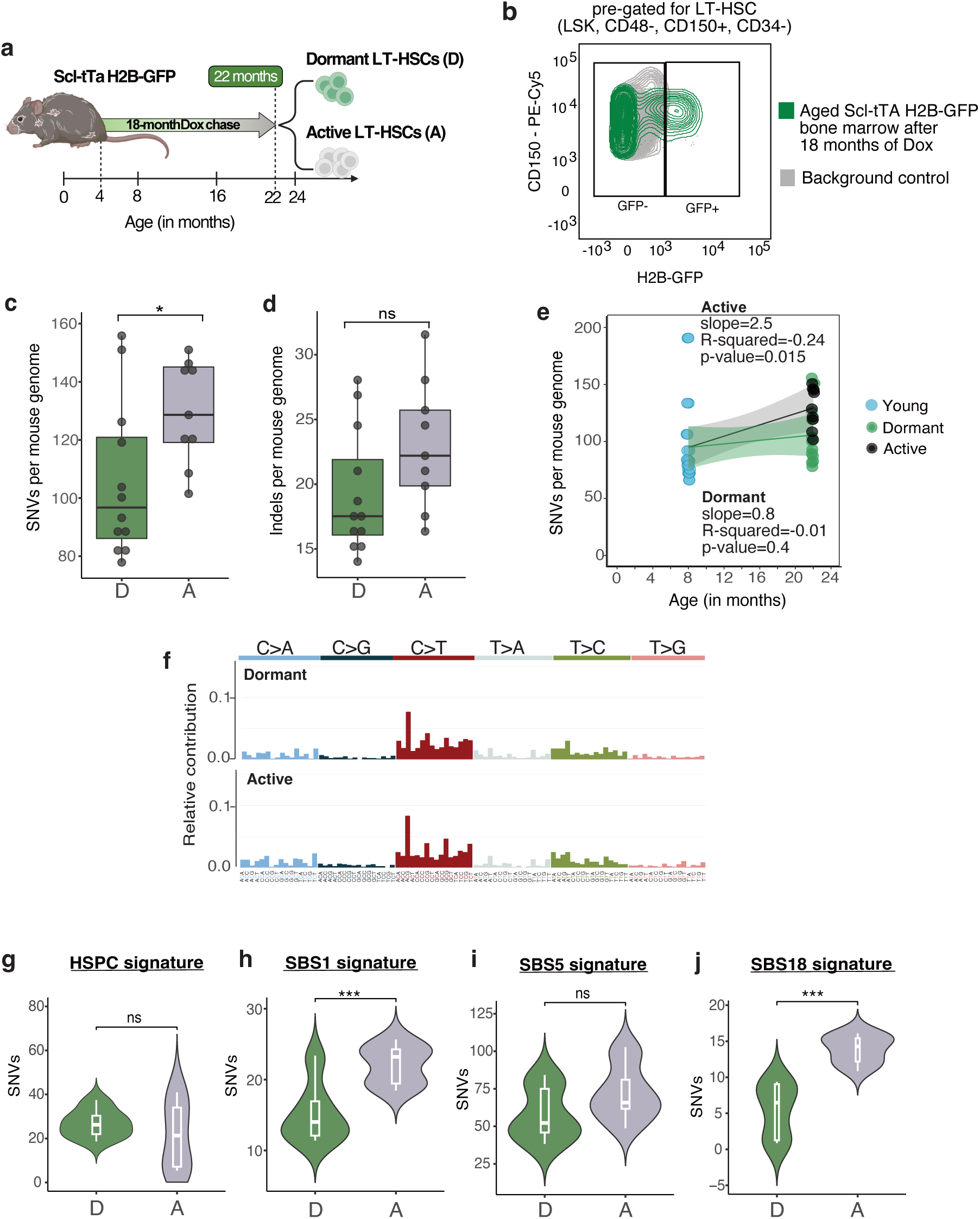
Dormancy protects LT-HSC from mutation acquisition during the lifetime of an experimental mouse. **a)** Schematic overview of the experimental design. Four-month-old Scl-tTA H2B-GFP mice were administered doxycycline (Dox) continuously via drinking water for 18 months, which comprises a chase period, where no new H2B-GFP is synthesized and existing H2B-GFP in chromatin is diluted by cell division. At 22 months of age, dormant and active LT-HSCs were isolated from the same aged donors (n=3 mice) and expanded *in vitro* prior to WGS. **b)** Exemplar flow cytometry gating strategy for the separation of dormant label-retaining and active non-label retaining LT-HSCs, based on GFP expression levels. The cut-off for GFP expression is defined by the highest level of fluorescence observed in LT-HSCs isolated from age-matched background control mice (H2B-GFP only), shown in grey. **c-d)** Absolute number of **c)** SNVs and **d)** indels, normalized per mouse genome for dormant (D) and active (A) LT-HSCs isolated from the same aged donor mice. **e)** Linear regression analysis of SNV burden as a function of age between young and old dormant LT-HSCs (green line) and between young and old active LT-HSCs (black line). Data for young LT-HSCs taken from Fig. 1). 95% CI is shaded. **f)** Trinucleotide SNV spectra of dormant and active LT-HSCs, merged per group. **g-j)** Absolute number of SNVs attributed to the stated fitted signature, normalized per mouse genome. Pairwise comparisons were assessed with Mann-Whitney U test. ns: p>0.05, *: p<0.05, ***: p<0.001. Each dot represents a single LT-HSC-derived colony., with n=12 (D) and n=9 (A). Box plots indicate median, inter-quartile range (IQR) with whiskers extending to 1.5x IQR.

### Sterlie inflammation accelerates accumulation of age-associated mutations in LT-HSCs

Dormant LT-HSCs can be provoked to enter into cell cycle under certain conditions of emergency haematopoiesis, such as in response to acute inflammatory stimulus or infection^41–45^. If genome stability is preserved by maintaining the dormant status, then one might predict that exposure to such stimuli might accelerate the rate at which LT-HSCs acquire mutations. In order to test this hypothesis, we employed a model of acute sterile inflammation which has been shown to induce the proliferation of dormant LT-HSC reserves and promote their contribution to all major mature blood cell lineages^46,47^. Thus, 2-month-old WT C57BL/6J mice were subject to three rounds of treatment with the TLR3 agonist polyinosinic-polycytidylic acid (pI:pC); each treatment round consisting of two injections per week for four weeks, interspersed with a 4-week recovery period (**Fig. 4a**). Our previous work has shown that this treatment regimen would result in a pronounced sampling bias if coupled with *in vitro* clonal expansion, since the LT-HSCs which proliferate in response to such a treatment regimen form smaller colonies and therefore would not generate sufficient genomic DNA to facilitate the necessary 30X coverage^46^. We therefore instead used NanoSeq^21^ which allowed us to directly enumerate the average mutation burden from the pool of ∼1,500 LT-HSCs we could isolate from each mouse without the need for *in vitro* expansion (**Fig. 4b**, **Fig S6a-c**). In line with our overarching hypothesis, LT-HSCs isolated from mice that had been exposed to the inflammatory regimen demonstrated an approximate 25% higher level of SNVs per genome compared to LT-HSCs from age-matched controls treated with phosphate buffered saline (PBS) (**Fig. 4c**), corresponding to the number of mutations that would be acquired following an additional 6-months of normal aging (**Fig. 1c**). There was also a trend towards an increased average number of indels per genome in the pI:pC-treated group, although the number of estimated indels per genome are low and Nanoseq is less accurate at detecting indels compared to SNVs^48^ (**Fig. 4d**). Importantly, mutation signature analysis revealed an analogous mutation spectrum to that observed during normal aging (**Fig 4e**, **Fig. 1f**), suggesting that the inflammatory stimulus did not promote a completely new form of DNA damage or mutagenic repair in LT-HSCs, but rather accelerated the processes which drive mutation acquisition during normal physiologic aging. Indeed, analysis of absolute levels of the HSPC SBS signature which accumulated in LT-HSC during normal aging, showed that it was enriched in LT-HSC from pI:pC treated mice compared to age-matched PBS-treated controls (**Fig. 4f-j**). This is in line with our previous data indicating that such an inflammatory insult can result in the *in vivo* induction of low-level DNA damage in LT-HSCs^35^, and can more generally accelerate irreversible aging of LT-HSCs^46^.

**Figure 4.**
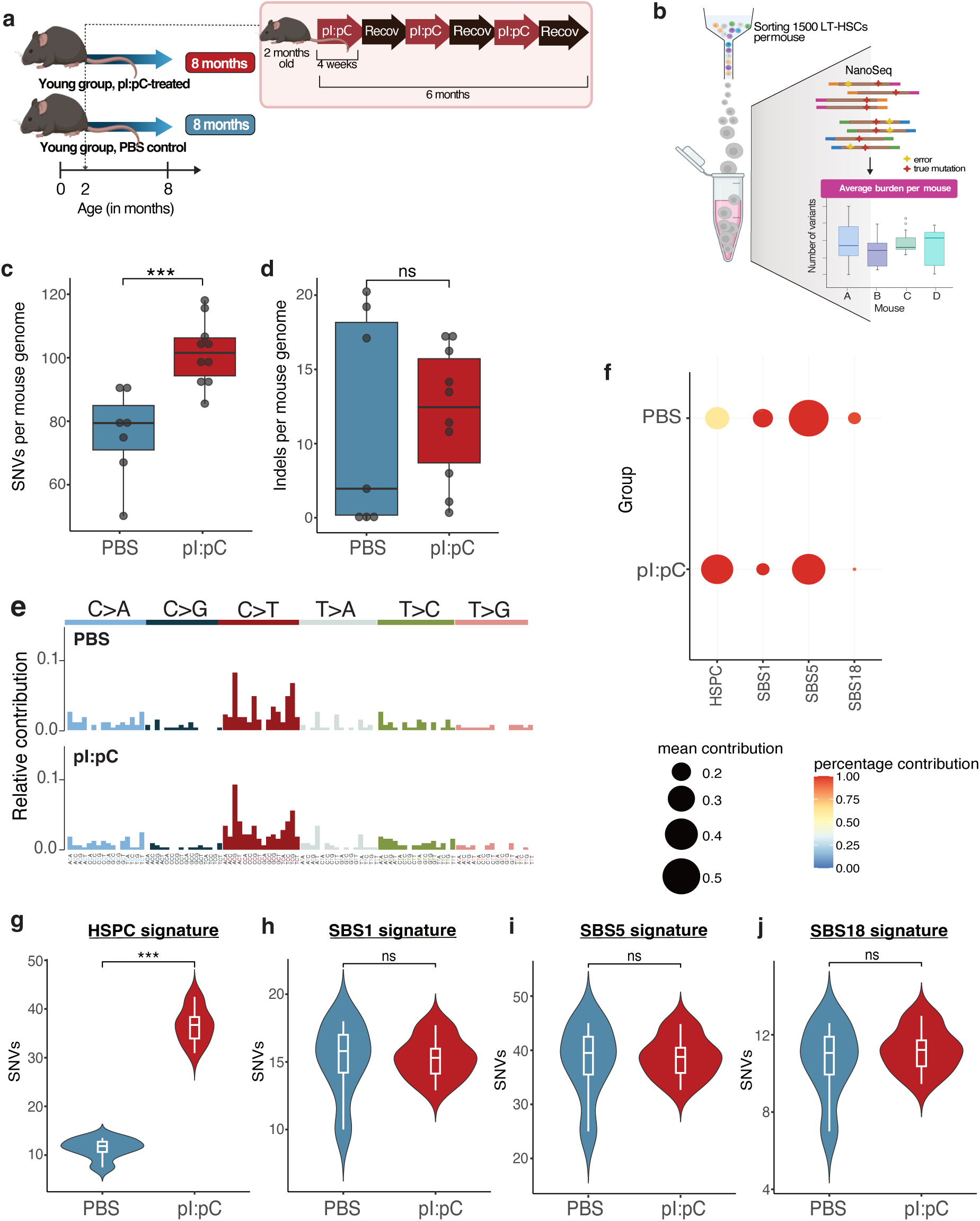
Inflammatory cues induce accelerated somatic mutagenesis in LT-HSCs *in vivo*. **a)** Schematic overview of the experimental design, where LT-HSCs were isolated from mice that had been repeatedly injected with either the TLR3 agonist pI:pC, or PBS. **b)** Schematic overview of mutation burden analysis in genomic DNA isolated from 1500 LT-HSCs per mouse, using NanoSeq. **c-d)** Absolute number of **c)** SNVs and **d)** indels, normalized per mouse genome. Each dot represents the average burden in LT-HSCs from an individual mouse. **e)** Trinucleotide SNV spectra of LT-HSCs from PBS-treated and pI:pC-treated mice. SNVs have been merged by group. **f)** Contribution of each fitted signature to the mutational profile of each treatment group. Circle size represents the mean signature contribution across 50 bootstrap iterations. Circle color indicates the percentage of bootstrap iterations in which the signature was detected. **g-j)** Absolute number of SNVs attributed to the indicated fitted signature, normalized per mouse genome. Two-group comparisons were assessed with Mann-Whitney U test. ns: p>0.05, ***: p<0.001. Boxplots indicate median, inter-quartile range (IQR) with whiskers extending to 1.5x IQR.

## Discussion

While the established dogma would firmly position programmed cell death as a critical gatekeeper that restricts the transition from DNA damage to mutation acquisition, our data indicates that genetic ablation of cell intrinsic apoptosis has no perceptible impact on mutation rate in HSCs *in vivo*. A likely explanation for this unexpected outcome is that the level of DNA damage that HSCs are exposed to during normal physiologic aging may be insufficient to cross the threshold necessary to trigger apoptosis, while classical studies demonstrating that apoptosis is a frequent outcome of the activation of the DDR typically employed genotoxic agonists such as chemotherapy or irradiation^11,22,23^. Nonetheless, it is clear that sufficient levels of DNA damage do occur during normal aging in order to facilitate a gradual accumulation of point mutations that equate to an average *in vivo* acquisition rate of 44 SNVs per year in murine LT-HSCs (**Fig 1d**). However, our study additionally documented the important finding that rate of mutation acquisition was highly heterogeneous across different LT-HSCs during aging, with dormant LT-HSCs accumulating barely any SNVs during the 18-month chase period, which equates to approximately 2/3 of the lifespan of C57BL/6J mice in our facility. We envisage that there are potentially two non-mutually exclusive explanations for the resistance of dormant LT-HSCs to mutagenesis during aging. Firstly, dormant LT-HSCs have gone through fewer rounds of DNA replication, meaning that they will have a lower lifetime exposure to replication stress as a potential source of DNA damage; will have lower accumulation of mutations due to replication by low-fidelity DNA polymerases; and have fewer opportunities for replication-associated repair processes to fix DNA lesions, DNA strand breaks, and slippage events as mutations in the genome. Secondly, dormancy might comprise a cellular state where LT-HSCs are exposed to lower levels of endogenous genotoxic metabolites, such as ROS, acetaldehyde and formaldehyde^49–51^. Indeed, dormant LT-HSCs are reported to exist in a metabolic state with decreased usage of oxidative phosphorylation and therefore lower potential to generate ROS through leakage in the electron transport chain, while the perisinusoidal niche that dormant LT-HSCs inhabit within the bone marrow cavity might limit extrinsic exposure to oxygen compared to their counterparts located at the arteriolar niche^34,52^. This would certainly correlate with our finding that the SBS18 signature is significantly reduced in aged dormant LT-HSCs compared to their active counterparts. Either way, our data corroborate the hypothesis that LT-HSC dormancy can act to uncouple the linear relationship between chronologic aging and the accumulation of DNA mutations and suggest that division-independent mutational mechanisms appear to play a minor role in the process of mutation acquisition within this dormant population.

An important implication of our study is that it challenges the concept that mutations are acquired in HSCs at a uniform rate with the passage of time during normal aging. Rather, it predicts that mutation rate can be modulated by extrinsic stimuli which impact on the state of dormancy, such as in the setting of stress hematopoiesis. Our data validates this prediction by showing that the level of SNVs in LT-HSCs is increased in mice exposed to sterile inflammation, and that the mutational spectrum is akin to that observed during normal aging, suggesting that this regimen resulted in an accelerated age-associated mutagenesis process as opposed to inducing a fundamentally different form of genomic instability. This has direct implications for the role of extrinsic factors such as infection and inflammation in modulating both the process of HSC aging and in the evolution of age-associated diseases that are the sequelae of somatic mutations within the stem cell compartment, such as cancer. Indeed, this conclusion is very much in line with the observation that past exposure to chronic infection and inflammation correlate with both the occurrence of age-associated pre-malignant clonal hematopoiesis^53^; and the incidence of acute myeloid leukemia and myelodysplastic syndrome^49^, both of which are age-associated myeloid malignant states in which the cell of origin is thought to be the LT-HSC. While recent studies suggest that such correlations might result from certain somatic mutations providing a selective growth advantage to the mutated HSC clone in the face of inflammatory challenge^17,54–56^, our data would suggest that inflammation may additionally contribute to this process by modulating the mutation burden within the HSC pool, therefore increasing the probability that one or more advantageous mutations are acquired in the first place. These findings have broad implications for our understanding of how environmental stimuli, which are not direct DNA damage agents, can nonetheless act as physiologic rheostats which modulate the rate of mutation acquisition in HSCs by transiently disrupting the protective dormant state, which otherwise preserves the genomic integrity of a sub-set of the HSC pool for extremely long periods of time.

## Acknowledgements

We would like to thank the DKFZ Flow Cytometry Core facility, the DKFZ center for preclinical research, the DKFZ Next Generation Sequencing Core facility and the Omics IT and Data Management Core facility, as well as the Sequencing Core Facility of the Wellcome Sanger Institute. R.B. and S.L. were supported by the DKFZ International Postdoctoral Fellowship Program; M.D. by the DKFZ-MOST German-Israeli Cooperation in Cancer Research Grant; and E.R.-C. by the German-Israeli Helmholtz International Research School “Cancer-TRAX”. This work was also supported by funding from the German Research Foundation (DFG) Collaborative Research Center CRC1709 (grant INST 35/1899-1 to M.D.M.); the Deutsche Jose Carreras Leukämiestiftung (grant R15/09 to M.D.M.); the Fritz Thyssen Stiftung (grant 10.16.1.023MN to M.D.M.); and the Forschungsinitiative Rheinland-Pfalz through the ReALity Excellence Program (to CI). We would also like to thank the Dietmar Hopp Foundation for their generous support.

## Author Contributions Statement

F.F., M.D., R.B., V.F., M.B., J.K., S.L., E.R.-C., K.A., I.G., M.B.S., A.M.M. and J.J. performed experiments. F.F., M.D., D.L., I.M., B.B., A.C. and M.D.M. designed experiments, provided technical expertise and interpreted data. F.F., M.D., V.F., A.B.-O., C.I. and A.C. performed bioinformatic analysis of sequencing data. F.F., A.C. and M.D.M. wrote the manuscript. I.M., B.B., A.C. and M.D.M. supervised the project.

---

**Figure S1.**
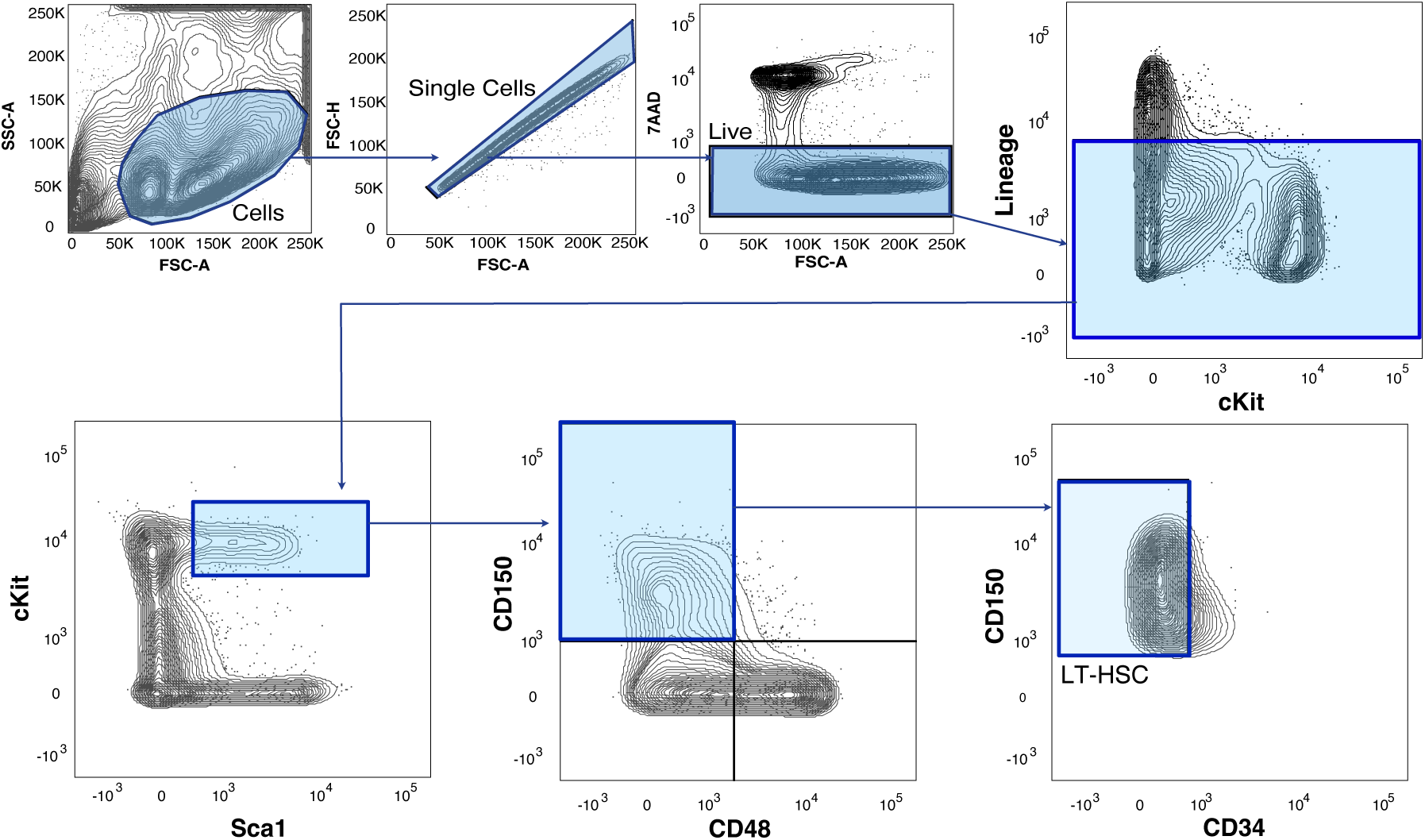
Flow cytometry gating strategy for isolation of LT-HSC from murine bone marrow. In this study, LT-HSC are defined as Lineage-; Sca-1+, c-Kit+, CD48-CD150+ CD34-.

**Figure S2.**
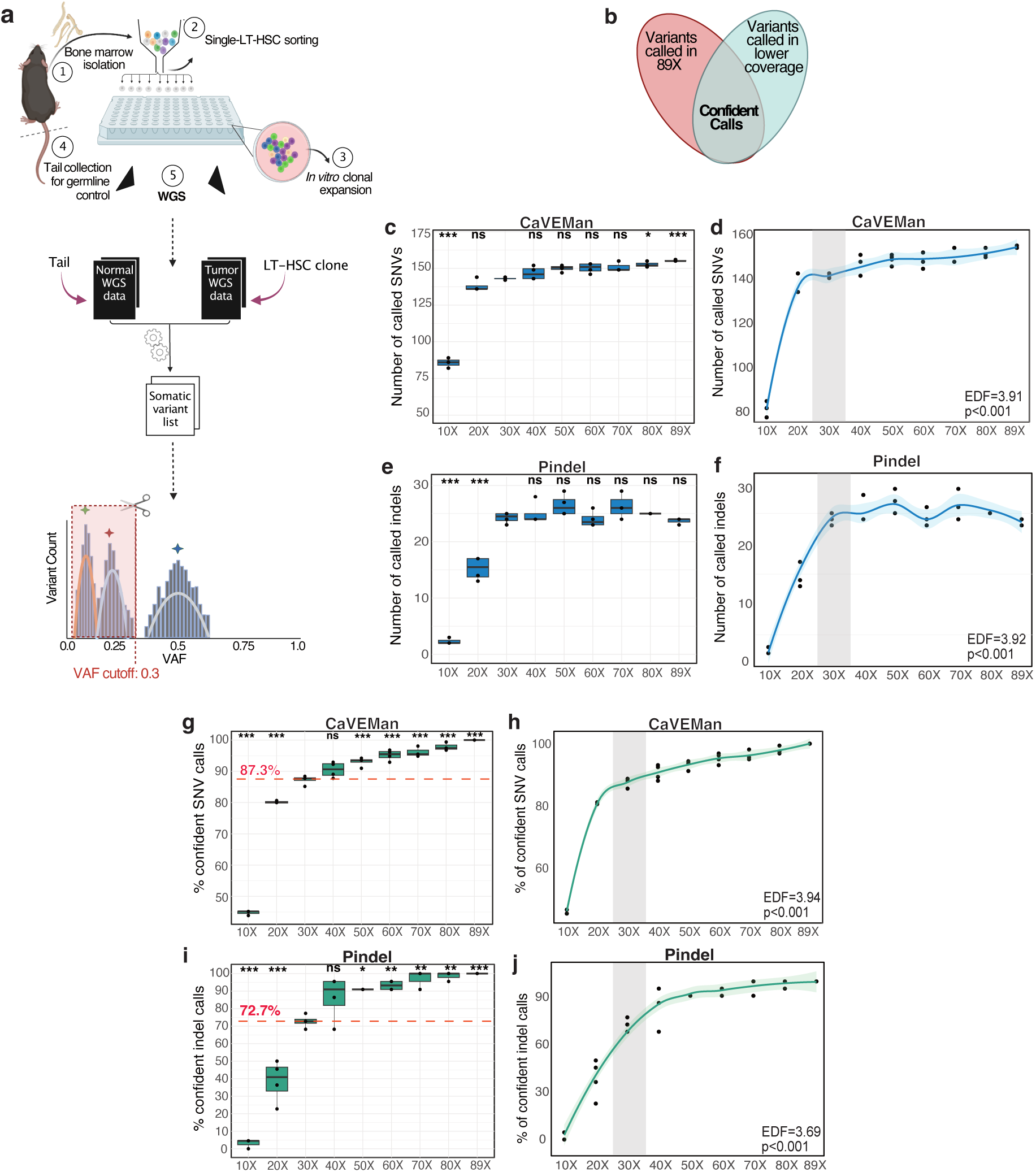
Quality control analysis of the effect of coverage on the accuracy and sensitivity of mutation detection in colonies derived from single LT-HSC clones. **a)** Experimental and analytical workflow for profiling single murine HSC genomes through clonal *in vitro* expansion of LT-HSC; followed by WGS of colonies and paired DNA from the tail; identification of somatic variants in the colonies; and filtering for mutations that were present in the original isolated LT-HSC using a VAF cutoff of 0.3. **b)** Definition of confident calls in benchmarking analysis based on overlap between variants called at the highest (89x) coverage versus those called in the down-sampled data sets. **c)** Number of SNVs detected at each down-sampled coverage level after quality filtering using CaVEMan. Boxplots indicate median, inter-quartile range (IQR) with whiskers extending to 1.5x IQR. **d)** Locally Estimated Scatterplot Smoothing (LOESS) curves using the data displayed in **c)**. The Effective Degrees of Freedom (EDF) value confirms the non-linear relationship between sequencing depth and caller performance (since EDF ≠ 1). Statistical significance was assessed by fitting a generalized additive model (GAM) to each curve. Blue bands represent 95% CI, and the gray rectangle marks the value at 30X coverage which was selected as the cut off value for subsequent experiments. **e)** Number of indels detected at each down-sampled coverage level after quality filtering using Pindel. Boxplots indicate median, inter-quartile range (IQR) with whiskers extending to 1.5x IQR. **f)** Locally Estimated Scatterplot Smoothing (LOESS) curves using the data displayed in **e). g)** Percentage of confident SNV calls identified in each coverage level, as defined in **b)**. Red line represents the mean caller performance at 30X coverage. **h)** Locally Estimated Scatterplot Smoothing (LOESS) curves using the data displayed in **g)**. Green shading represents 95% CI, and the gray rectangle marks the 30X coverage value. **i)** Percentage of confident indel calls identified in each coverage level. Red line represents the mean caller performance at 30X coverage. **j)** Locally Estimated Scatterplot Smoothing (LOESS) curves using the data displayed in **i)**. Green shading represents 95% CI, and the gray rectangle marks the 30X coverage value. Statistical annotations in **c)** and **d)** reflect comparisons of each coverage group against the 30X group, assessed using one-way ANOVA, followed by Tukey’s HSD correction for multiple comparisons.

**Figure S3.**
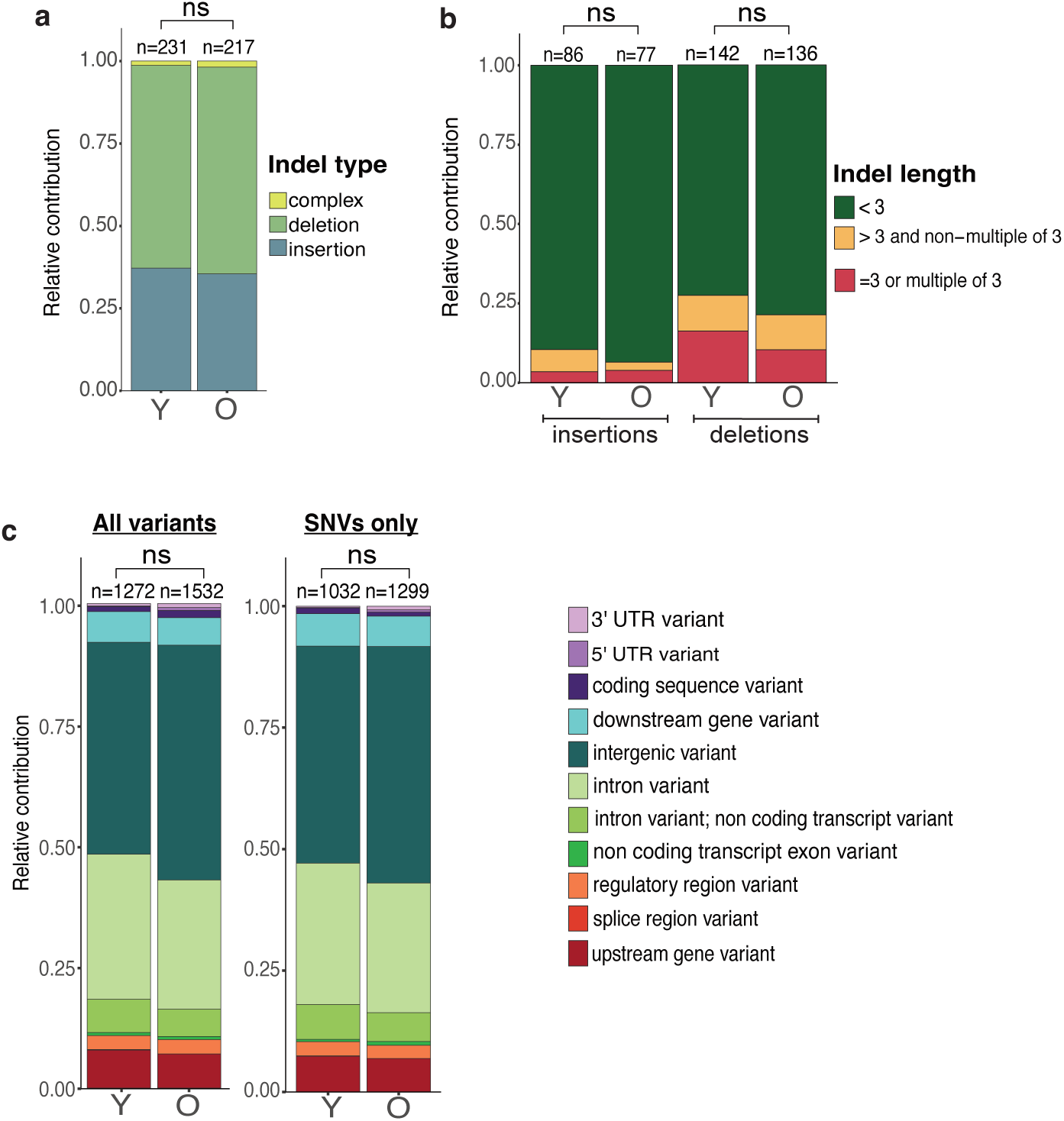
Analysis of mutation type and genomic location in young versus old LT-HSCs. **a)** Categorization of indels identified in young and old LT-HSCs. “Complex” indels refer to events involving both an insertion and a deletion at the same locus. **b)** Sub-segregation of indels in young and old LT-HSCs according to length of region inserted or deleted. **c)** Genomic annotation of all the variants (left) and of the SNVs only (right) detected in young and old LT-HSCs. Proportional differences were assessed with chi-square tests. ns: p>0.05. The number of variants per group is indicated on the corresponding stacked bars. Y: young, O: old.

**Figure S4.**
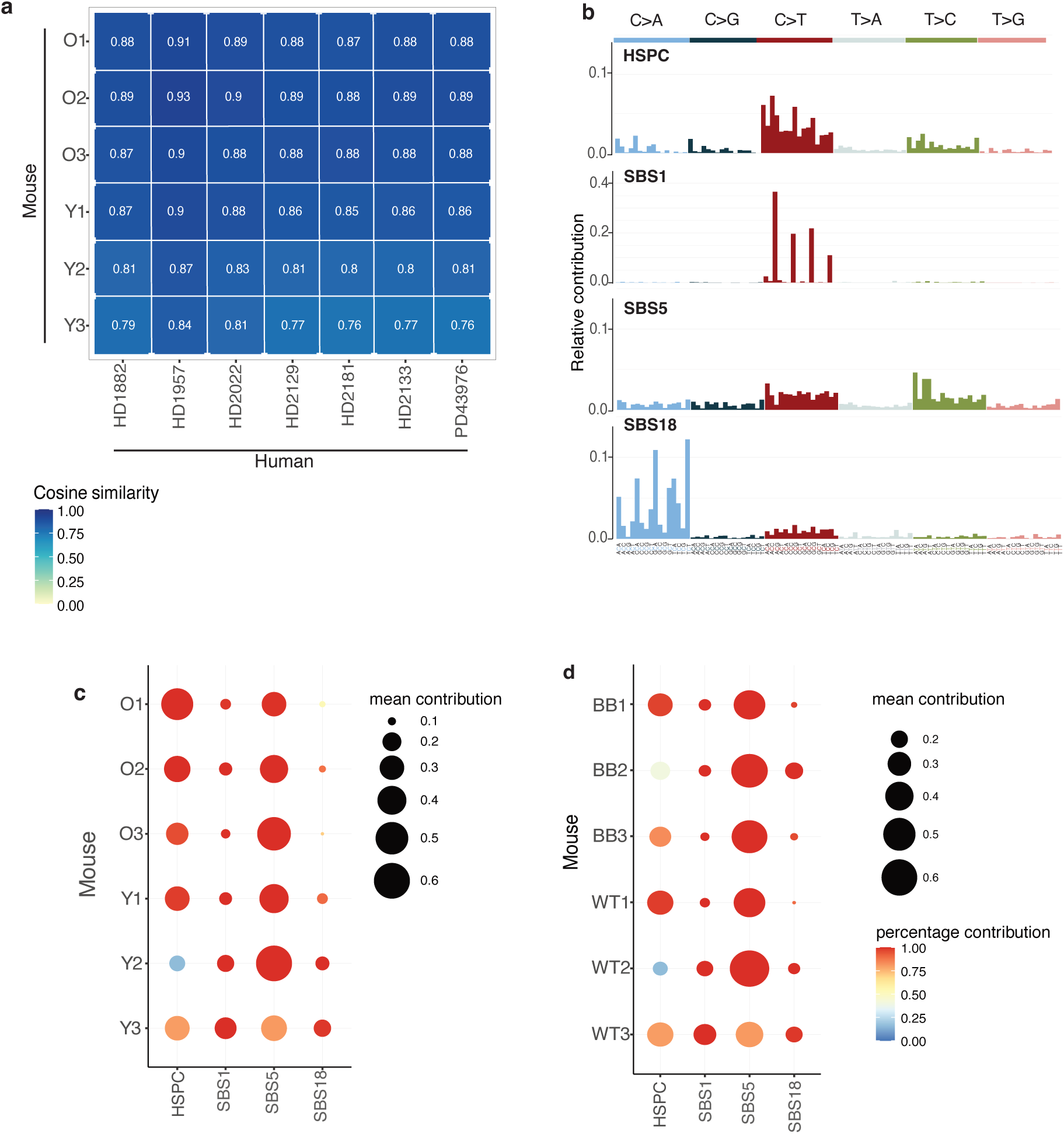
Comparative analysis of trinucleotide mutation signatures in LT-HSCs between young and old, or wild-type and Bak^-/-^ Bax^Δ/Δ^ experimental groups. **a)** Cosine similarities between the trinucleotide mutational profiles of young and old mice and the healthy adult human donors from Abascal et al.^18^ **b)** Trinucleotide patterns of the four known signatures that were identified in our dataset through signature fitting. **c-d)** Contribution of each fitted signature to the mutational profile of individual mice. Circle size represents the mean signature contribution across 150 bootstrap iterations. Circle colour indicates the percentage of bootstrap iterations in which the signature was detected. **c**) Young (Y) versus old (O). **d**) Wild-type (WT) versus Bak^-/-^ Bax^Δ/Δ^ (BB). The mouse IDs BB3 corresponds to data generated from LT-HSCs that were isolated from three different, but co-housed mice, that were merged and collectively analysed as a single mouse for more robust signature analysis.

**Figure S5.**
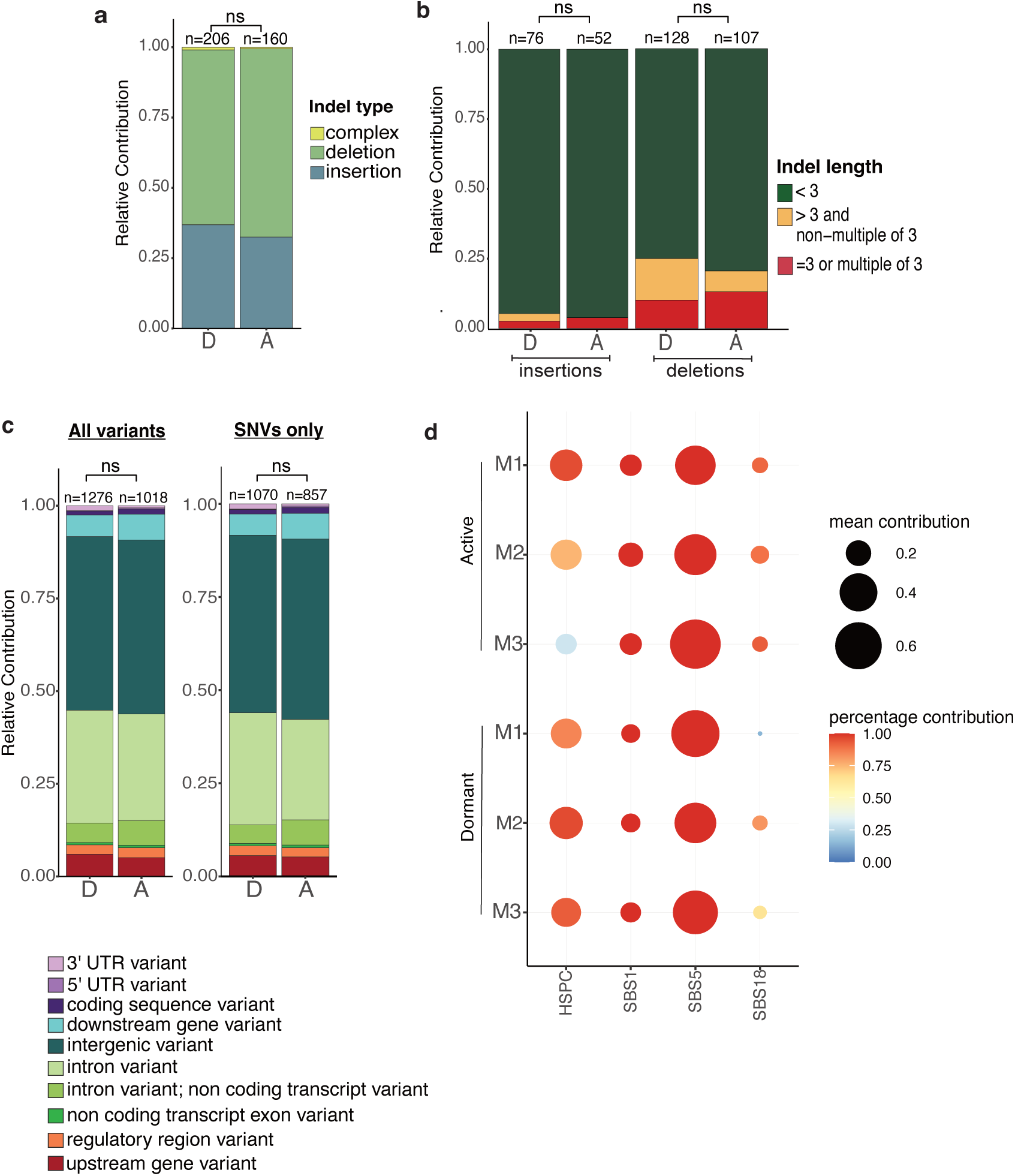
Analysis of mutation type, genomic location and mutational spectrum in dormant versus active LT-HSCs isolated from old mice. **a)** Sub-segregation of the different indel types identified in dormant and active HSC genomes. **b)** Proportions of different indel lengths detected in dormant and active HSC genomes. **c)** Genomic annotation of all the variants (left) and the SNVs only (right) detected in dormant and active HSCs. **d)** Contribution of each fitted signature to the mutational profile of individual mice. Circle size represents the mean signature contribution across 150 bootstrap iterations. Circle colour indicates the percentage of bootstrap iterations in which the signature was detected. M1-M3 indicates the experimental mouse from which the dormant and active LT-HSCs were isolated from. Pairwise comparisons were performed using the Mann-Whitney U test. Differences in the proportions were assessed with a chi-square test. ns: p>0.05. *: p<0.05. The number of variants per group is indicated on the corresponding stacked bars. D: dormant, A: active

**Figure S6.**
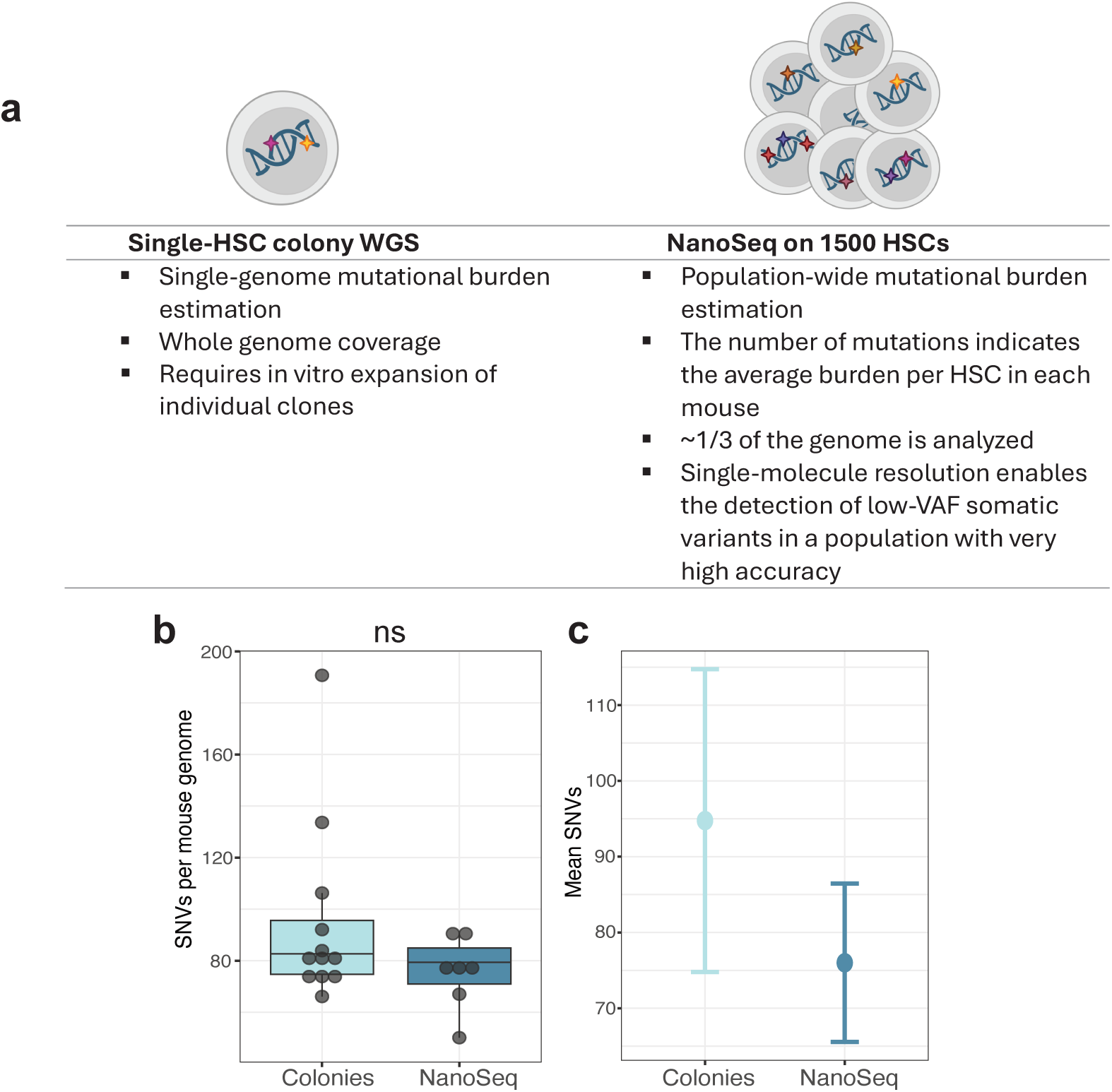
Comparison of mutation analysis using whole genome sequencing of a single LT-HSC-derived colony versus Nano-seq of a population of purified LT-HSCs. **a)** Summary of the fundamental differences between the single-HSC clonal WGS performed on DNA from colonies, versus the NanoSeq approach performed on DNA isolated from a polyclonal (1500 HSCs) mini-bulk population. **b)** Left: Comparison of single-HSC mutational burden estimated by WGS of HSC-derived colonies versus the average single-cell mutational burden inferred from NanoSeq, performed on HSC mini-bulks. Each dot represents analysis of the clonal progeny from an individual cell in the colony-derived WGS group, whereas each dot represents average mutation burden of LT-HSCs isolated from a single mouse in the NanoSeq group. For this comparison, colony-derived WGS data from three young (8-month-old) mice were used (from Figure 1), and NanoSeq data were obtained from seven PBS-treated 8-month-old mice. Statistical significance between groups was assessed using the Mann-Whitney U test. ns: p>0.05. For “Colonies”, each dot represents the SNV burden of an individual LT-HSC-derived clone, while in “NanoSeq” it represents the average SNV burden of the interrogated bulk LT-HSC population per mouse. Boxplots indicate median, inter-quartile range (IQR) with whiskers extending to 1.5x IQR. **c)** Mean SNV burden per cell as measured by the two approaches. Error bars mark the 95% confidence intervals (CI).

## Methods

### Mouse models

All animal experiments were approved by the Animal Care and Use Committees of the German Regierungspräsidium Karlsruhe für Tierschutz und Arzneimittelüberwachung (Karlsruhe, Germany) under the TVAs G-117/17 and G-284/18. All mouse models were bred and maintained at the DKFZ Central Animal Laboratory under specific pathogen-free (SPF) conditions. Mice were housed in individually ventilated cages (IVC) in same-sex groups of up to five individuals. The same samples served as the young control group for Figure 1 and Figure 2. To induce the homozygous deletion of the floxed Bax gene in Scl-CreERT2 *Bak*^-/-^ *Bax*^fl/fl^ HSCs, mice received five intraperitoneal (i.p.) injections of 200 µL tamoxifen (10 mg/ml; Sigma-Aldrich) dissolved in 10% ethanol (Supelco) and 90% sunflower seed oil (Sigma-Aldrich). For the label-retention assay, 16-week-old *Scl-tTA H2B-GFP* mice were administered doxycycline (2 mg/ml; Sigma-Aldrich) with 20 mg/ml sucrose through their drinking water. WT mice (C57BL/6J) received 3 rounds of intermittent i.p. injections of pI:pC (5 mg/kg; InvivoGen), prepared in sterile phosphate-buffered saline (PBS) (Sigma) according to the manufacturer’s guidelines. Each treatment round consisted of two injections per week for four weeks, followed by a four-week recovery period. Mice were 2 months old at the beginning of the treatment and approximately 8 months old at its conclusion.

### Bone marrow isolation

Tibiae, femora, ilia, vertebrae, and sterna were harvested after removing adherent flesh with a scalpel. The bones were crushed in Iscove’s Modified Dulbecco’s Medium (IMDM), and the resulting cell suspension was filtered through 40 μm cell strainers (Greiner Bio-One) to eliminate bone fragments, centrifuged, and resuspended in PBS - 2% FBS (Gibco). Bone marrow counts were measured with a ScilVet-abc Plus+ veterinary counter (Scil GmbH).

### Tail collection

To capture the genomic variations of the germline for each mouse, a 2-3 cm segment of the tail was cut post-mortem. For the subsequent DNA extraction, 2 mg of tissue were processed by adhering to the DNeasy Blood and Tissue Kit (Qiagen) protocol.

### Single LT-HSC sorting

Bone marrow cell suspension was subjected to depletion of mature blood cell lineages by incubating with a mix of rat anti-mouse biotin-conjugated lineage markers (4.2 µg/mL CD5, 4.2 µg/mL CD8a, 2.4 µg/mL CD11b, 2.8 µg/mL B220, 2.4 µg/mL Gr-1, 2.6 µg/mL Ter-119) for 40 min at 4°C. After incubation, cells were washed once with PBS + 2% FBS, spun down at 350 g at 4°C for 5 min, resuspended in 800 µL PBS + 2% FBS, and mixed with 800 µL Biotin Binder Dynabeads (Thermo Fisher), which were previously washed (2 washes with PBS + 2% FBS). Beads were added at a concentration of 1 mL beads /1x10^8^ cells. Cells-beads mix was incubated for 45 min at 4°C with constant rotation. Subsequently, lineage-positive cells were depleted using a magnetic particle concentrator (Dynal MPC-6, Invitrogen), and the resulting LSK-enriched fraction was washed once with PBS + 2% FBS and stained with the panel of antibodies indicated in SupplementaryTable 1a and 1b for 30 minutes at 4°C in the dark. After the incubation, stained cells were washed once with PBS + 2% FBS, resuspended in a final concentration of 2 mL PBS + 2% FBS, and filtered through a 40 µm cell strainer FACS tube before the sort. Single LT-HSCs were sorted using Aria I or II sorters (BD) and collected into round-bottom 96-well plates (Cellstar) filled with 200 µL/well of serum-free expansion medium (SFEM; StemSpan) (**Fig. S1**).

### LT-HSC-mini bulk sorting for Nanoseq

A mini bulk of 1500 LT-HSCs from the stained bone marrow suspension (Supplementary Table 1a) was sorted per mouse (n=7 for PBS-treated and n=10 for pI:pC-treated mice) into a single well of a skirted 96-well plate. For direct cell lysis, the PicoPure DNA Extraction Kit (ThermoFischer) was used. Mini bulks were sorted into 20 ul of Reconstitution Buffer supplemented with Proteinase K, following the manufacturer’s instructions. Plates were then sealed and centrifuged at 1500 rpm for 30 sec at 4°C, followed by incubation in a thermal cycler for 3 hours at 65oC and 10 minutes at 95°C.

### *In vitro* clonal expansion of single LT-HSCs

*In vitro* clonal expansion of single HSCs was performed as previously described by Bogeska et al., 2022^46^. At the end of the incubation period, wells were examined under a light microscope to assess clonogenic efficiency, calculated as the number of wells with colony formation of any size divided by the total number of wells. Approximately 30 colonies per mouse were selected and reserved for WGS.

### DNA extraction from LT-HSC colonies

Genomic DNA was extracted from colonies using the QIAmp DNA mini kit (Qiagen) following the manufacturer’s guidelines. The only deviation from the standard protocol was the elution step: DNA was eluted in 30 ul of nuclease-free water instead of the recommended 200 ul, to obtain more concentrated DNA samples. DNA concentration was measured with a Qubit 3.0 Fluorometer (Life Technologies) using the Qubit dsDNA High Sensitivity Assay Kit (Life Technologies).

### WGS library preparation and sequencing

WGS libraries were prepared using the TruSeq Nano DNA Low Throughput Library Prep Kit (Illumina) with adherence to the manufacturer’s instructions. Although the protocol recommends a minimum DNA input of 100 ng, most colonies yielded less DNA. To accommodate lower input amounts, the sonication duration and the number of PCR amplification cycles were optimized as described in Supplementary Table 2. Library quality was assessed using a Bioanalyzer 2100 with High Sensitivity DNA chips (Agilent). Paired-end 150 bp WGS was performed using either a Hi-Seq X10, NovaSeq 6000 or NovaSeq X series sequencer. 2-12 colonies per mouse, from 3-5 mice per group, together with the respective tails, were eventually sequenced at 30X minimum coverage. Samples with WGS data failing the quality control standards defined as duplication rates >25%, coverage <30X, and lack of clonal behaviour based on VAFs were excluded from the study.

### Benchmarking dataset preparation

DNA was isolated from a large HSC colony using the DNAeasy Blood and Tissue Kit (Qiagen). WGS libraries were prepared following the tagmentation-based whole-genome bisulfite sequencing protocol^46^, excluding the bisulfite conversion step. Up to 5 ng of DNA was used as input per library, and the amount of Tn5 transposase was adjusted accordingly. To achieve deep sequencing coverage, five individually prepared libraries and one quadruplex library were generated for both the colony and the matched germline control. Each library was sequenced in a dedicated lane of the Illumina HiSeq V4 platform using 125 bp paired-end reads.

### NanoSeq library preparation

Whole-genome NanoSeq libraries were prepared as previously described^18^ from sorted HSC mini-bulks. Genomic DNA was fragmented by restriction enzyme digestion according to the original NanoSeq protocol. Following end repair and adapter ligation, A-tailing was performed in the presence of dideoxynucleotides to suppress extension from single-strand nicks and minimise artefactual transfer of errors between strands. Libraries were then qPCR-bottlenecked to optimise duplex recovery. Matched tail DNA was processed in parallel using the same library preparation protocol, but without the bottleneck step, to enable genotyping of inherited germline variation.

### Nanoseq sequencing

NanoSeq libraries were sequenced as 150-bp paired-end reads on an Illumina NovaSeq X platform (25B flow cells). HSC mini-bulk libraries were loaded at 0.3 fmol and sequenced to a mean depth of 30X per sample. Matched tail libraries were loaded at 3-5 fmol and sequenced in parallel to a mean depth of 15X per sample for germline SNP genotyping.

### Read alignment and genotyping

Sequencing reads were mapped to the mouse reference genome (GRCm38) using the Burrows-Wheeler Aligner^57^ (BWA; version 0.7.15). Duplicate reads were identified and marked with Sambamba^58^ (version 0.6.5). For sequenced clones derived from Scl-CreERT2 *Bak*^-/-^ *Bax*^fl/fl^ mice, alignments were manually inspected using the Integrative Genomics Viewer^59^ (IGV, version 2.13.2) to confirm effective Cre-mediated deletion.

### SNV calling

For the HSC clones, SNVs were identified using CaVEMan^15^ (version 1.16.0). This algorithm detects putative somatic base substitutions by comparing sequencing data from a test sample, its matched normal (germline), and the reference genome (here: GRCm38, mm10). In this study, each aligned HSC clone sequence was compared to the corresponding tail tissue from the same mouse to subtract the germline variants and retain only those present exclusively in the clones. Repetitive genomic regions, including centromeric and simple repeats, were excluded from the analysis. Additionally, a panel of normal (PON) mouse samples sequenced under similar conditions was used to filter out common genomic regions prone to errors, such as mapping artefacts or polymorphisms. The resulting Variant Call Format (VCF) files were further filtered as listed below:

– PASS filter: Only variants marked with the “PASS” flag by CaVEMan, indicating they met all internal quality thresholds, were retained.
– Sex chromosome filter: Sex chromosomes were removed since both male and female mice were included in the dataset and were unevenly distributed across the different experimental groups.
– Coverage filter: Variants found in regions with sequencing depth below 10X and above 100X were excluded to avoid low-confidence calls and potential mapping artefacts.
– Alignment filters: Variants of a read-adjusted median alignment score (ASRD) less than 0.94 were removed. Additionally, variants supported by reads containing soft-clipped bases (Clipped Read Position Model, CLPM) were also filtered out (i.e., CLPM must be =0).
– VAF filter: Heterozygous somatic mutations in the originally sorted LT-HSC were expected to exhibit a VAF around 0.5. Variants with VAF below 0.3 were filtered out to eliminate *in vitro*-acquired mutations. Samples lacking this characteristic distribution were considered non-clonal and excluded from downstream analysis.

For NanoSeq, aligned reads were processed using the NanoSeq bioinformatic pipeline^18^, adapted for mouse samples. Somatic SNVs were called from duplex consensus data using the NanoSeq workflow, applying the standard NanoSeq quality filters. To account for the absence of equivalent mouse resources for known SNPs and recurrent artefactual sites, we generated a mouse-specific mask from the sequenced tail samples. This comprised sites supported by at least one variant read in at least two mice, which were classified as recurrent noisy sites, and sites within a single mouse sample with a variant allele fraction greater than or equal to 0.2, or number of reads greater than or equal to 10, which were classified as candidate germline SNPs. Masked positions were excluded from somatic variant calling in all samples and from calculation of callable genome space.

### Indel calling

Small insertions and deletions were identified using Pindel^16^ (version 3.9.0). The maximum detectable insertion size was limited by read length, approximately 130 bp in this study. Like CaVEMan, Pindel detects somatic indels by comparing aligned sequences from each LT-HSC clone with the corresponding tail tissue alignments from the same mouse. Centromeric regions, simple repeats, and other unreliable mouse genomic regions were excluded from the analysis. Post-calling, additional filtering steps were applied. Variants that failed Pindel’s internal quality filters (FF004, FF005, FF006, FF012, FF015, and FF016) were discarded. Additionally, a minimum Phred quality score of 20 was required. All remaining calls were subjected to the same downstream filters applied to the SNV dataset.

For Nanoseq, small indels were identified from duplex consensus data using the indel-calling workflow distributed with the NanoSeq codebase, applying the same core quality-control and positional filters described above.

### SV calling

Larger SVs were identified using Delly^60^ (version 1.1.5). In the initial step, Delly excludes highly repetitive regions, such as telomeric and centromeric sequences. Each colony was compared to its matched germline control, and candidate somatic SVs were additionally screened against a PON constructed from all germline samples in the dataset. Delly’s internal filters discard SVs supported by fewer than 3 reads (from either paired-end or split-end evidence) or with mapping quality <20. For inter-chromosomal translocations, a minimum of 5 supporting reads was required. Following Delly’s built-in filters, additional post-calling filters were applied, including those based on coverage, VAF, and exclusion of sex chromosomes.

### Benchmarking analysis

Coverage downsampling was performed on a deeply sequenced LT-HSC colony (initial coverage 89X) and its matched tail (initial coverage 97X) using downsampleSAM from Picard^61^ (version 1.61). Downsampled fractions were calculated to approximate coverage levels in 10X increments: 10X, 20X, 30X, 40X, 50X, 60X, 70X, 80X, and 89X. These target depths were achieved by adjusting the “PROBABILITY” parameter in the downsampling tool. Each coverage level was generated in triplicate using different seeds (1, 2, and 3), resulting in 3 alignment files (BAM) per coverage point. Each depth-matched LT-HSC clone and tail alignment pair was used for variant calling with CaVEMan and Pindel following the approach explained above. For the benchmarking analysis, the coverage filter <100X was deprecated. At full coverage (89X), the original merged HSC alignment was compared to each of the three downsampled tail alignments. Confident calls were defined as those consistently detected across all three comparisons. To confirm the relationship between sequencing coverage and number of variants called and percentage of confident calls we applied locally estimated scatterplot smoothing (LOESS) using the “loess()” function in R. The optimal LOESS span parameter was selected via leave-one-out cross-validation (LOOCV) by minimizing mean squared error across a range of span values. To statistically assess the significance of the observed trends, we also fit generalized additive models (GAMs) using the “gam()” function from the mgcv^62^ (version 1.9-3) R package.

### Callable genome estimation and variant normalization

To facilitate comparisons across samples and groups from the LT-HSC-colony-derived data, variant counts were normalized based on the size of the callable genome for each sample. Per-base coverage was assessed with Mosdepth^63^ (version 0.3.3), categorizing regions into coverage bins: 0-10X, 10- 100X, and >100X. Regions falling in the former and the latter coverage range in both colony and tail samples were excluded from downstream analysis using BEDTools^64^ (version 2.29.2). From the retained 10-100X regions, additional exclusions were applied to remove areas ignored by the variant callers, regions with variants that did not meet the alignment or quality criteria, and recurring problematic regions identified in the PON. The callable genome size was thus defined as the sum of reliable, well-covered autosomal regions per sample. Final variant counts were normalized using the formula: (number of variants) x (reference autosomal genome size)/(callable autosomal genome size).

For Nanoseq-derived data, genome-wide somatic SNV burden was estimated from filtered duplex consensus calls and normalised to callable genome space. For each sample, the somatic mutation rate per base pair was calculated by dividing the number of filtered somatic SNVs by the number of callable duplex bases after exclusion of masked sites. Per-cell somatic SNV burden was then estimated by multiplying this rate by the size of the callable diploid mouse genome.

### Variant annotation

All the filtered variants were annotated using the Ensembl Variant Effect Predictor^65^ (VEP, version 102.0). The primary consequence per variant was determined using the flag “–flag_pick”, which selects the most severe consequence based on VEP’s default prioritization criteria.

### Mutational profile and signature analysis

To construct SNV mutational profiles and extract the underlying signatures, the MutationalPatterns^66^ R package (version 3.12.0) was used. Prior to analysis, VCF files containing only the filtered SNVs per LT-HSC colony were concatenated to generate one VCF file per mouse. For the Nanoseq results, the filtered calls were concatenated per treatment group. Assuming similar mutagenic processes within age and activity groups, samples were analyzed in three sets: 1) young and old, 2) dormant and active cells, and 3) apoptosis-deficient cells. Due to the low mutational burdens, known signature fitting was performed via bootstrapping, using the “fit_to_signatures_bootstrapped” function. For each mouse, the average contribution of each signature was calculated by averaging across all bootstrap iterations. Absolute SNV counts per signature were then estimated by multiplying these average contributions by the normalized SNV counts per cell from the corresponding mouse. For the comparison of mutational profiles between mouse and human, we used the colony-derived data from healthy adult donors from Abascal et al. 2021^21^.

### Data visualization and statistics

Data analysis, visualization, and further statistical testing were performed using R (version 4.3.0). Sorting data was analyzed and visualized using FlowJo v10. Statistical comparisons across two groups were conducted with Mann-Whitney U test. Linear regression was conducted using the “lm” function. Differences in variant annotation frequencies between groups were evaluated using chi-squared tests.

## Data availability

The data for this study have been deposited in the European Nucleotide Archive (ENA) at EMBL-ENI under accession numbers PRJEB82854 (https://www.ebi.ac.uk/ena/browser/view/PRJEB82854) and ERP192755 (https://www.ebi.ac.uk/ena/browser/view/ERP192755).

## Code availability

The code that was generated in this study is available in Zenodo with the DOI: https://doi.org/10.5281/zenodo.19979487

---

**Supplementary Table 1a.** Antibody panel for single LT-HSC and mini-bulk (Nanoseq) sorting using WT and Bak-/- Bax^Δ/Δ^ mice.

| Antigen | Fluorophore | Supplier; Catalog number |
| --- | --- | --- |
| CD4 | eFluor450 | eBioscience; 48-0041-82 |
| CD8a | eFluor450 | eBioscience; 48-0081-82 |
| CD11b | eFluor450 | eBioscience; 48-0112-82 |
| B220 | eFluor450 | eBioscience; 48-0452-82 |
| Gr1 | eFluor450 | eBioscience; 48-5931-82 |
| Ter119 | eFluor450 | eBioscience; 48-4921-82 |
| Streptavidin | eFluor450 | eBioscience; 48-4317-82 |
| cKit (CD117) | BV711 | Biolegend; 105835 |
| Sca-1 | APC-Cy7 | BD Biosciences; 560654 |
| CD150 | PE-Cy5 | Biolegend; 115912 |
| CD48 | PE-Cy7 | Biolegend; 103424 |
| CD34 | FITC | eBioscience; 11-0341-82 |
| Viability dye | 7AAD | Invitrogen; A1310 |

**Supplementary Table 1b.** Antibody panel for single LT-HSC sorting using Scl-tTA H2B-GFP mice.

| Antigen | Fluorophore | Supplier; Catalog number |
| --- | --- | --- |
| CD4 | PE-Cy7 | eBioscience; 48-0041-82 |
| CD8a | PE-Cy7 | eBioscience; 48-0081-82 |
| CD11b | PE-Cy7 | eBioscience; 48-0112-82 |
| B220 | PE-Cy7 | eBioscience; 48-0452-82 |
| Gr1 | PE-Cy7 | eBioscience; 48-5931-82 |
| Ter119 | PE-Cy7 | eBioscience; 48-4921-82 |
| Streptavidin | PE-Cy7 | eBioscience; 25-4317-82 |
| cKit (CD117) | APC | eBioscience; 17-1171-83 |
| Sca-1 | APC-Cy7 | BD Biosciences; 560654 |
| CD150 | PE-Cy5 | Biolegend; 115912 |
| CD48 | PE | eBioscience; 12-0481 |
| CD34 | eFluor450 | eBioscience; 48-0341-82 |
| Viability dye | 7AAD | Invitrogen; A1310 |

**Supplementary Table 2.**
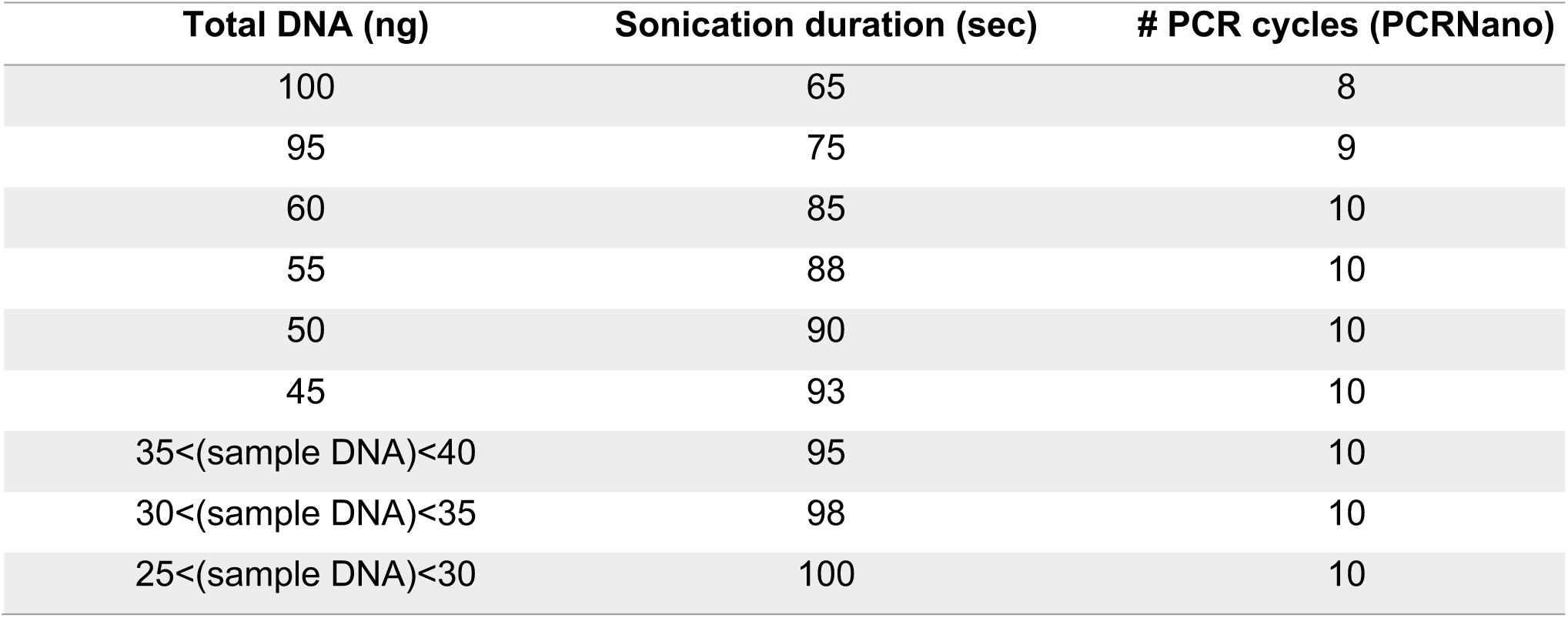
Sonication duration and number of PCR cycles used during WGS library preparation, adjusted according to input DNA amount.

## References

1. López-Otín, C., Blasco, M. A., Partridge, L., Serrano, M. & Kroemer, G. Hallmarks of aging: An expanding universe. Cell 186, 243–278 (2023).

2. Hanahan, D. Hallmarks of Cancer: New Dimensions. Cancer Discovery 12, 31–46 (2022).

3. Schumacher, B., Pothof, J., Vijg, J. & Hoeijmakers, J. H. J. The central role of DNA damage in the ageing process. Nature 592, 695–703 (2021).

4. Panier, S., Wang, S. & Schumacher, B. Genome Instability and DNA Repair in Somatic and Reproductive Aging. Annu. Rev. Pathol. Mech. Dis. 19, 261–290 (2024).

5. Ciccia, A. & Elledge, S. J. The DNA Damage Response: Making It Safe to Play with Knives. Molecular Cell 40, 179–204 (2010).

6. d’Adda Di Fagagna, F. Living on a break: cellular senescence as a DNA-damage response. Nat Rev Cancer 8, 512–522 (2008).

7. Vijg, J. From DNA damage to mutations: All roads lead to aging. Ageing Research Reviews 68, 101316 (2021).

8. Biechonski, S., Yassin, M. & Milyavsky, M. DNA-damage response in hematopoietic stem cells: an evolutionary trade-off between blood regeneration and leukemia suppression. Carcinogenesis 38, 367–377 (2017).

9. Rossi, D. J., Jamieson, C. H. M. & Weissman, I. L. Stems Cells and the Pathways to Aging and Cancer. Cell 132, 681–696 (2008).

10. Lee, N., Spears, M. E., Carlisle, A. E. & Kim, D. Endogenous toxic metabolites and implications in cancer therapy. Oncogene 39, 5709–5720 (2020).

11. Huang, R. & Zhou, P.-K. DNA damage repair: historical perspectives, mechanistic pathways and clinical translation for targeted cancer therapy. Sig Transduct Target Ther 6, 254 (2021).

12. Nowsheen, S. & Yang, E. S. The intersection between DNA damage response and cell death pathways. Exp Oncol 34, 243–254 (2012).

13. Lee-Six, H. et al. Population dynamics of normal human blood inferred from somatic mutations. Nature 561, 473–478 (2018).

14. Alioto, T. S. et al. A comprehensive assessment of somatic mutation detection in cancer using whole-genome sequencing. Nat Commun 6, 10001 (2015).

15. Jones, D., et al. cgpCaVEManWrapper: Simple Execution of CaVEMan in Order to Detect Somatic Single Nucleotide Variants in NGS Data. CP in Bioinformatics 56, (2016).

16. Ye, K., Schulz, M. H., Long, Q., Apweiler, R. & Ning, Z. Pindel: a pattern growth approach to detect break points of large deletions and medium sized insertions from paired-end short reads. Bioinformatics 25, 2865–2871 (2009).

17. Kapadia, C. D. et al. Clonal dynamics and somatic evolution of haematopoiesis in mouse. Nature 641, 681–689 (2025).

18. Alexandrov, L. et al. Signatures of mutational processes in human cancer. Nature 500, 415–421 (2013).

19. Osorio, F. G. et al. Somatic Mutations Reveal Lineage Relationships and Age-Related Mutagenesis in Human Hematopoiesis. Cell Reports 25, 2308–2316.e4 (2018).

20. De Kanter, J. K. et al. Antiviral treatment causes a unique mutational signature in cancers of transplantation recipients. Cell Stem Cell 28, 1726–1739.e6 (2021).

21. Abascal, F. et al. Somatic mutation landscapes at single-molecule resolution. Nature 593, 405–410 (2021).

22. Matt, S. & Hofmann, T. G. The DNA damage-induced cell death response: a roadmap to kill cancer cells. Cell. Mol. Life Sci. 73, 2829–2850 (2016).

23. Roos, W. P. & Kaina, B. DNA damage-induced cell death by apoptosis. Trends in Molecular Medicine 12, 440–450 (2006).

24. Opferman, J. T. et al. Obligate Role of Anti-Apoptotic MCL-1 in the Survival of Hematopoietic Stem Cells. Science 307, 1101–1104 (2005).

25. Qing, Y., Wang, Z., Bunting, K. D. & Gerson, S. L. Bcl2 overexpression rescues the hematopoietic stem cell defects in Ku70-deficient mice by restoration of quiescence. Blood 123, 1002–1011 (2014).

26. Milyavsky, M. et al. A Distinctive DNA Damage Response in Human Hematopoietic Stem Cells Reveals an Apoptosis-Independent Role for p53 in Self-Renewal. Cell Stem Cell 7, 186–197 (2010).

27. Yu, H. et al. Deletion of Puma protects hematopoietic stem cells and confers long-term survival in response to high-dose γ-irradiation. Blood 115, 3472–3480 (2010).

28. Moehrle, B. M. et al. Stem Cell-Specific Mechanisms Ensure Genomic Fidelity within HSCs and upon Aging of HSCs. Cell Reports 13, 2412–2424 (2015).

29. Takeuchi, O. et al. Essential role of BAX,BAK in B cell homeostasis and prevention of autoimmune disease. Proc. Natl. Acad. Sci. U.S.A. 102, 11272–11277 (2005).

30. Wei, M. C. et al. Proapoptotic BAX and BAK: A Requisite Gateway to Mitochondrial Dysfunction and Death. Science 292, 727–730 (2001).

31. Lindsten, T. et al. The Combined Functions of Proapoptotic Bcl-2 Family Members Bak and Bax Are Essential for Normal Development of Multiple Tissues. Molecular Cell 6, 1389–1399 (2000).

32. Göthert, J. R. et al. In vivo fate-tracing studies using the Scl stem cell enhancer: embryonic hematopoietic stem cells significantly contribute to adult hematopoiesis. Blood 105, 2724–2732 (2005).

33. Bernitz, J. M., Kim, H. S., MacArthur, B., Sieburg, H. & Moore, K. Hematopoietic Stem Cells Count and Remember Self-Renewal Divisions. Cell 167, 1296–1309.e10 (2016).

34. Saçma, M. et al. Haematopoietic stem cells in perisinusoidal niches are protected from ageing. Nat Cell Biol 21, 1309–1320 (2019).

35. Walter, D. et al. Exit from dormancy provokes DNA-damage-induced attrition in haematopoietic stem cells. Nature 520, 549–552 (2015).

36. Mohrin, M. et al. Hematopoietic Stem Cell Quiescence Promotes Error-Prone DNA Repair and Mutagenesis. Cell Stem Cell 7, 174–185 (2010).

37. Beerman, I., Seita, J., Inlay, M. A., Weissman, I. L. & Rossi, D. J. Quiescent Hematopoietic Stem Cells Accumulate DNA Damage during Aging that Is Repaired upon Entry into Cell Cycle. Cell Stem Cell 15, 37–50 (2014).

38. Wilson, A. et al. Hematopoietic Stem Cells Reversibly Switch from Dormancy to Self-Renewal during Homeostasis and Repair. Cell 135, 1118–1129 (2008).

39. Blokzijl, F. et al. Tissue-specific mutation accumulation in human adult stem cells during life. Nature 538, 260–264 (2016).

40. Kucab, J. E. et al. A Compendium of Mutational Signatures of Environmental Agents. Cell 177, 821–836.e16 (2019).

41. Essers, M. A. G. et al. IFNα activates dormant haematopoietic stem cells in vivo. Nature 458, 904–908 (2009).

42. Baldridge, M. T., King, K. Y., Boles, N. C., Weksberg, D. C. & Goodell, M. A. Quiescent haematopoietic stem cells are activated by IFN-γ in response to chronic infection. Nature 465, 793–797 (2010).

43. Takizawa, H. et al. Pathogen-Induced TLR4-TRIF Innate Immune Signaling in Hematopoietic Stem Cells Promotes Proliferation but Reduces Competitive Fitness. Cell Stem Cell 21, 225–240.e5 (2017).

44. Takizawa, H., Regoes, R. R., Boddupalli, C. S., Bonhoeffer, S. & Manz, M. G. Dynamic variation in cycling of hematopoietic stem cells in steady state and inflammation. Journal of Experimental Medicine 208, 273–284 (2011).

45. Haltalli, M. L. R. et al. Manipulating niche composition limits damage to haematopoietic stem cells during Plasmodium infection. Nat Cell Biol 22, 1399–1410 (2020).

46. Bogeska, R. et al. Inflammatory exposure drives long-lived impairment of hematopoietic stem cell self-renewal activity and accelerated aging. Cell Stem Cell 29, 1273–1284.e8 (2022).

47. Munz, C. M. et al. Regeneration following blood loss and acute inflammation proceeds without contribution of primitive HSCs. Blood blood.2022018996 (2023) doi:10.1182/blood.2022018996.

48. Cheng, Y. et al. Improved Mutation Detection in Duplex Sequencing Data with Sample-Specific Error Profiles. Preprint at 10.1101/2025.07.13.664565 (2025).

49. Garaycoechea, J. I. et al. Alcohol and endogenous aldehydes damage chromosomes and mutate stem cells. Nature 553, 171–177 (2018).

50. Garaycoechea, J. I. et al. Genotoxic consequences of endogenous aldehydes on mouse haematopoietic stem cell function. Nature 489, 571–575 (2012).

51. Pontel, L. B. et al. Endogenous Formaldehyde Is a Hematopoietic Stem Cell Genotoxin and Metabolic Carcinogen. Molecular Cell 60, 177–188 (2015).

52. Jang, Y.-Y. & Sharkis, S. J. A low level of reactive oxygen species selects for primitive hematopoietic stem cells that may reside in the low-oxygenic niche. Blood 110, 3056–3063 (2007).

53. Lapite, A. et al. Association of infection frequency and incident clonal hematopoiesis of indeterminant potential. Experimental Hematology 158, 105420 (2026).

54. Jakobsen, N. A. et al. Selective advantage of mutant stem cells in human clonal hematopoiesis is associated with attenuated response to inflammation and aging. Cell Stem Cell 31, 1127–1144.e17 (2024).

55. Rodriguez-Meira, A. et al. Single-cell multi-omics identifies chronic inflammation as a driver of TP53-mutant leukemic evolution. Nat Genet 55, 1531–1541 (2023).

56. Hormaechea-Agulla, D. et al. Chronic infection drives Dnmt3a-loss-of-function clonal hematopoiesis via IFNγ signaling. Cell Stem Cell 28, 1428–1442.e6 (2021).

## Methods-only references

57. Li, H. & Durbin, R. Fast and accurate short read alignment with Burrows–Wheeler transform. Bioinformatics 25, 1754–1760 (2009).

58. Tarasov, A., Vilella, A. J., Cuppen, E., Nijman, I. J. & Prins, P. Sambamba: fast processing of NGS alignment formats. Bioinformatics 31, 2032–2034 (2015).

59. Robinson, J. T. et al. Integrative genomics viewer. Nat Biotechnol 29, 24–26 (2011).

60. Rausch, T. et al. DELLY: structural variant discovery by integrated paired-end and split-read analysis. Bioinformatics 28, i333–i339 (2012).

61. Picard Toolkit. (2019). Broad Institute, GitHub Repository. https://broadinstitute.github.io/picard/; Broad Institute

62. Wood, S. N. Fast Stable Restricted Maximum Likelihood and Marginal Likelihood Estimation of Semiparametric Generalized Linear Models. Journal of the Royal Statistical Society Series B: Statistical Methodology 73, 3–36 (2011).

63. Pedersen, B. S. & Quinlan, A. R. Mosdepth: quick coverage calculation for genomes and exomes. Bioinformatics 34, 867–868 (2018).

64. Quinlan, A. R. & Hall, I. M. BEDTools: a flexible suite of utilities for comparing genomic features. Bioinformatics 26, 841–842 (2010).

65. McLaren, W. et al. The Ensembl Variant Effect Predictor. Genome Biol 17, 122 (2016).

66. Manders, F. et al. MutationalPatterns: the one stop shop for the analysis of mutational processes. BMC Genomics 23, 134 (2022).

